# Phylogenetic modeling of regulatory element turnover based on epigenomic data

**DOI:** 10.1101/773614

**Authors:** Noah Dukler, Yi-Fei Huang, Adam Siepel

## Abstract

Evolutionary changes in gene expression are often driven by gains and losses of cis-regulatory elements (CREs). The dynamics of CRE evolution can be examined using multi-species epigenomic data, but so far such analyses have generally been descriptive and model-free. Here, we introduce a probabilistic modeling framework for the evolution of CREs that operates directly on raw chromatin immunoprecipitation and sequencing (ChIP-seq) data and fully considers the phylogenetic relationships among species. Our framework includes a phylogenetic hidden Markov model, called epiPhyloHMM, for identifying the locations of multiply aligned CREs, and a combined phylogenetic and generalized linear model, called phyloGLM, for accounting for the influence of a rich set of genomic features in describing their evolutionary dynamics. We apply these methods to previously published ChIP-seq data for the H3K4me3 and H3K27ac histone modifications in liver tissue from nine mammals. We find that enhancers are gained and lost during mammalian evolution at about twice the rate of promoters, and that turnover rates are negatively correlated with DNA sequence conservation, expression level, and tissue breadth, and positively correlated with distance from the transcription start site, consistent with previous findings. In addition, we find that the predicted dosage sensitivity of target genes positively correlates with DNA sequence constraint in CREs but not with turnover rates, perhaps owing to differences in the effect sizes of the relevant mutations. Altogether, our probabilistic modeling framework enables a variety of powerful new analyses.

## Introduction

It is now well established that the evolution of form and function is often driven by mutations in cis-regulatory elements (CREs), particularly in multicellular eukaryotes having complex programs for regulating gene expression^1–4^. In humans, patterns of genetic polymorphism, patterns of interspecies divergence, and the results of genome-wide association studies all indicate that a majority of phenotype- or fitness-influencing nucleotides fall in noncoding sequences and likely function in gene regulation^5–10^. While the mutations that underly regulatory evolution sometimes have subtle effects on, say, protein-DNA binding or chromatin accessibility, in many of the best-known cases, they instead alter gene expression through the gain or loss in activity of a whole CRE^11–13^. A number of lines of evidence indicate that this gain-and-loss process—sometimes called “turnover”—occurs at substantial rates over evolutionary time^14–22^. Indeed, the evolutionary dynamics of this process appear to play out over considerably shorter time periods than those for other critical functional elements, such as protein-coding genes, microRNAs, or long noncoding RNAs^23, 24^.

There have been numerous attempts to model the evolutionary dynamics of CRE turnover at the level of the primary DNA sequence^14, 15, 25–27^. However, characterizing this process at the sequence level is fundamentally challenging owing to limitations in the inference of regulatory function from the DNA sequence alone. During the past 15 years, new technologies for collecting high-throughput epigenomic data—such as chromatin immunoprecipitation and sequencing (ChIP-seq) data for transcription factors or histone modifications—have provided a path forward, by more directly indicating similarities and differences across species in molecular phenotypes that are closely related to cis-regulatory activity. A considerable number of comparative epigenomic studies have now been carried out in a variety of organisms, including studies based on transcription factor binding^17, 18, 28–31^, particular histone modifications^32–36^, chromatin accessibility or chromatin contacts^37–39^, DNA methylation^40^, and nascent transcription^24^ (partially reviewed in ref. ^41^). Among other findings, these studies have confirmed generally rapid rates of CRE gain and loss, and demonstrated that turnover rates are substantially higher in enhancers than in promoters, that depth of conservation correlates with various measures of functional impact, and that the evolutionary stability of gene expression correlates with the complexity and conservation of the local CRE architecture. However, with rare exceptions^40, 42^ (see **Discussion**), the available comparative epigenomic data sets have been analyzed using heuristic, model-free methods that do not consider the phylogenetic relationships of the species under study or the uncertainty in epigenomic data.

In this article, we introduce new model-based inference methods that address these deficiencies by fully accounting for the species phylogeny as well as the relationship between element activity and raw ChIP-seq read counts. Our methods can also account for correlations of turnover rates with local features along the genome sequence—such as gene expression patterns across tissues, distance to the transcription start site, or DNA sequence conservation—and they are efficient enough to be applied to genome-wide data sets. As a proof of concept, we apply these methods to previously published ChIP-seq data for the H3K4me3 and H3K27ac histone modifications in liver tissue across a phylogeny of nine mammals^35^. As described in detail below, we confirm several previous findings regarding relative rates of turnover in enhancers and promoters, and correlations with gene expression patterns and local regulatory architecture. In addition, we examine differences between patterns of constraint at the DNA sequence and CRE turnover levels, and find evidence suggesting that they reflects differences in the effect sizes of the relevant mutations.

## Results

### General approach

Our approach for analyzing multi-species epigenomic data consists of three major stages (Fig. 1A). First, we carry out a series of preprocessing steps to summarize the ChIP-seq read counts for each species in a common coordinate system (based on version hg38 of the human reference genome), excluding genomic regions where we could not establish clear one-to-one orthology based on genomic synteny (see **Methods** for details). Second, we apply a newly developed probabilistic inference method, called epiPhyloHMM, to identify “active” regions based on the ChIP-seq read counts, working in the common coordinate system. At this stage, an “active” region is one containing CREs in any one or more species. This method accounts for the phylogenetic gain and loss process, as well as noise in the ChIP-seq data, at the same time as it predicts the locations of the elements. Third, we apply a new probabilistic modeling program, called phyloGLM, to describe the process of phylogenetic gain and loss in more detail, within the “active” regions identified by epiPhyloHMM. PhyloGLM conditions on a rich set of genomic features, capturing their correlations with local rates of gain and loss.

**Figure 1:**
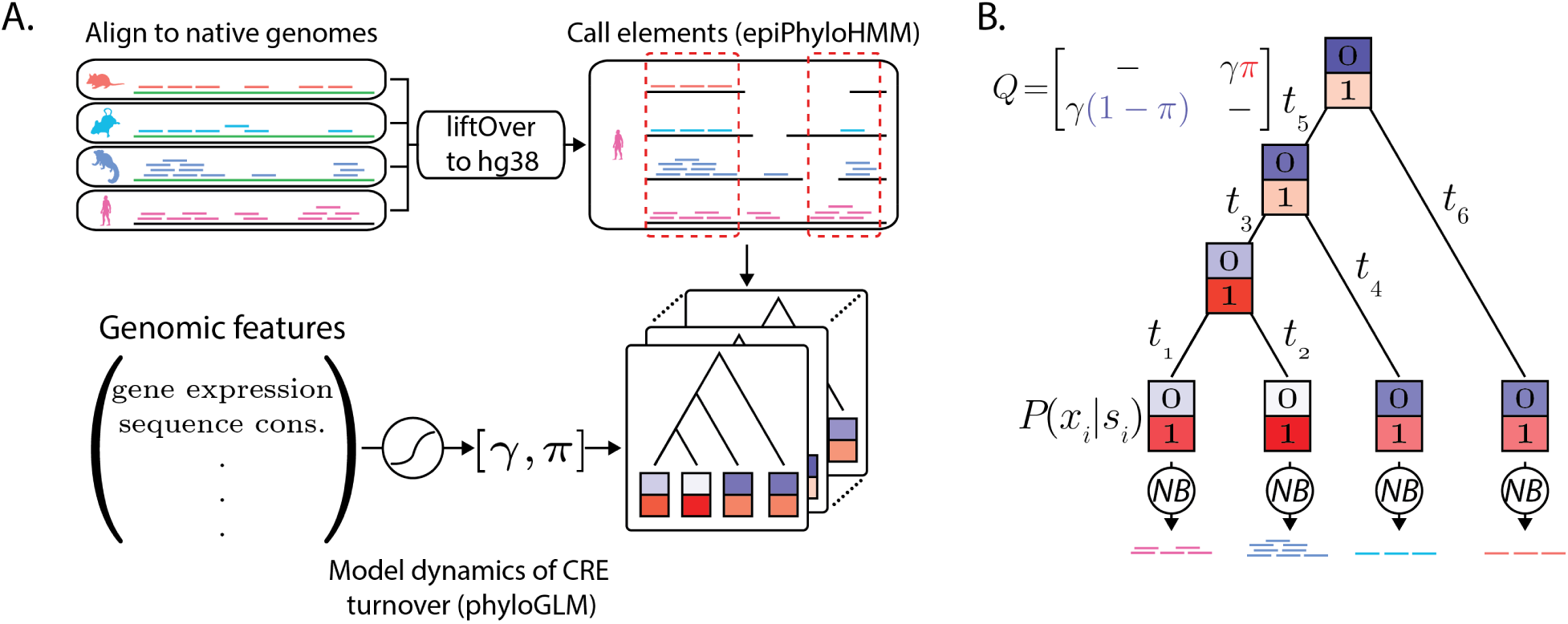
Illustration of modeling framework. (A) ChIP-seq data is aligned separately to the reference genome for each species then converted to the coordinate system of the human (hg38) genome using the liftOver program (**Methods**). Only regions of apparent one-to-one orthology are considered, based on synteny. Cis-regulatory elements (CREs) that are active in one or more species are then identified using epiPhyloHMM. Finally, the dynamics of CRE turnover within these elements are modeled using phyloGLM, which accounts for the associations between various genomic features and local rates of gain and loss. (B) Both epiPhyloHMM and phyloGLM use a core phylogenetic model in which the presence (*s_i_* = 1) and absence (*s_i_* = 0) of CREs is allowed to change in a branch-length-dependent manner along a fixed phylogeny, according to a continuous-time Markov model. The model is defined by an instantaneous rate matrix *Q* (dashes indicate values required for rows to sum to zero). The probabilities of the ChIP-seq read counts at the tips of the tree (*x_i_*) given *s_i_* are modeled using negative binomial (NB) distributions. The color intensities for the “0” and “1” boxes are proportional to the corresponding conditional likelihoods. π: stationary frequency of CRE presence; γ: gain/loss rate; *t_i_*: length of branch *i*.

### Shared phylogenetic model

The epiPhyloHMM and phyloGLM programs both make use of the same core probabilistic model for the gain and loss of CREs along the branches of a phylogeny. Moreover, in both cases, this model also describes the generation of read counts that are reflective of CRE presence or absence at the tips of the tree. Thus, it serves as a generative model for multi-species read counts that can be fitted to the raw data by maximum likelihood (Fig. 1B). In this article, we focus on read counts from ChIP-seq experiments, but the model can easily be extended to other data types, such as those arising from DNase-seq, ATAC-seq, or PRO-seq experiments.

The phylogenetic component of the model is a straightforward presence/absence model (with state variables *s* ∈ {0, 1}) for CREs along the branches of a phylogeny. It assumes a tree topology is given together with nonnegative real-valued branch lengths. In practice, the tree and branch lengths can be obtained from the literature or estimated from sequence data (see **Methods**). The stochastic process for gains and losses is defined, in the usual manner, by a continuous-time Markov model with an instantaneous rate matrix *Q*, from which branch-length dependent turnover (gain/loss) probabilities can be obtained as *P*(*t*) = exp(*Qt*) for each branch-length *t* (see ref. ^43^). The model has two free parameters: the stationary probability of CRE presence (*π*) and a single turnover rate parameter (*γ*), which together define a reversible rate matrix *Q* (Fig. 1B). Given data at the leaves of the tree, phylogenetic inference with this model can be accomplished using Felsenstein’s pruning algorithm^44, 45^ (see **Methods**).

Unlike with standard phylogenetic models for DNA sequences, however, the observed data here consists of epigenomic (typically ChIP-seq) read counts, which provide only an approximate indication of whether or not an active CRE exists in each species. We accommodated the uncertainty in read counts by borrowing from the literature on statistical peak calling for ChIP-seq data^46^. In particular, we described both the probability of the observed read counts *x_i_* in species *i* given an active CRE in that species, *P*(*x_i_* | *s_i_* = 1), and the probability of the observed read counts given no active CRE, *P*(*x_i_* | *s_i_* = 0), using negative binomial distributions (Fig. 1B). Moreover, for the “active” model, we used a mixture of three negative binomial distributions to accommodate peaks of various heights (see **Methods**)^47, 48^. We also adapted these emission distributions to accommodate missing data due to alignment gaps (Methods). Altogether, this modeling approach allows us to perform multi-species peak calling and phylogenetic inference simultaneously, accounting for uncertainty in both the locations of present day CREs along the genome and their presence/absence over evolutionary time.

### epiPhyloHMM: Prediction of multi-species CREs from epigenomic data

To address the problem of predicting “active” CREs, we made use of the framework of phylogenetic hidden Markov models, or phylo-HMMs^49, 50^. Phylo-HMMs are hidden Markov models whose hidden states are associated with phylogenetic models, which in turn, define distributions over columns in multiply aligned sequences of observations. Phylo-HMMs are sometimes called “space-time” models^51^ because they describe stochastic processes in both a spatial dimension, along the genome sequence, and a temporal dimension, along the branches of a phylogeny. In this case, the temporal (phylogenetic) models describe distributions over aligned ChIP-seq readcounts from multiple species, as described in the previous section. The spatial (hidden Markov) model, in turn, is designed to allow the identification of CREs with various patterns of presence/absence at the tips of the tree.

This hidden Markov model consists of a single “inactive” state and a set of states representing each possible presence/absence pattern (Fig. 2A). Assuming most of the genome will be inactive, the transition model is sparse, with each active state being accessible only from the inactive state, and not from other active states (see ref. ^25^ for a similar approach). It is completely defined by two free parameters, ρ_0_ and ρ_1_ (Fig. 2A and Methods). In practice, we constrain the complexity of the model by including states only for presence/absence patterns that are achievable by at most three gain/loss events along the branches of the phylogeny (see **Methods**). The free parameters of both the phylogenetic model (*π, γ*) and the HMM (*ρ*_0_, *ρ*_1_) are fitted to aligned epigenomic data by maximum likelihood, and then active elements are called in the standard way, using the Viterbi algorithm (Fig. 2B). The method generally performs well on simulated data (Supplemental Methods & Supplemental Figs. S1–S5).

**Figure 2:**
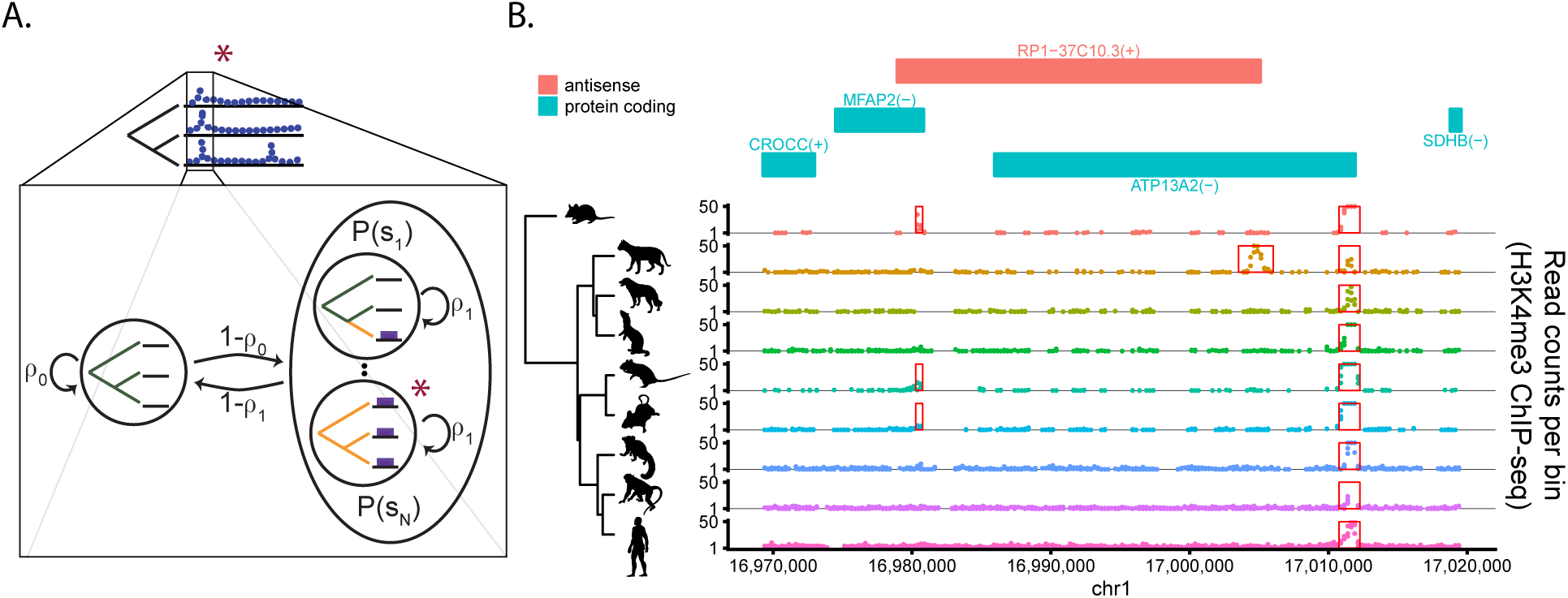
Prediction of multi-species cis-regulatory elements (CREs) using epiPhyloHMM. (A) State-transition diagram for the phylogenetic hidden Markov model. Each state represents a different combination of active and inactive elements in the observed species, corresponding to a different gain/loss scenario along the branches of the tree (*gold*: active; *black*: inactive). The fully inactive (background) state is shown on the *left* and the states representing various possible presence/absence patterns for active elements are grouped on the *right*. The red star indicates the state associated with the focal site in the cartoon data at top. Notice that each active state is accessible only from the inactive state, leading to a sparse transition matrix for the hidden Markov model. (B) Example of predictions obtained by applying epiPhyloHMM to H3K4me3 ChIP-seq data from ref. ^35^ in a region along human chromosome 1. Shown are the nine-species phylogeny (*left*) and the corresponding ChIP-seq read counts along the chromosome (*right*), with predicted elements highlighted in red boxes. For reference, annotated protein-coding genes and an antisense transcription unit are also shown (*top*).

### Application of epiPhyloHMM to histone-modification data for nine mammals

We applied epiPhyloHMM to recently published H3K4me3 and H3K27ac ChIP-seq data for liver tissue from mammals^35^, preprocessing and aligning the data as outlined above (see also Methods). We used data for nine of the 20 species examined in ref. ^35^, prioritizing ones with high-quality genome assemblies and alignments. This analysis produced an average of ∼16,000 and ∼47,000 elements per species for the H3K4me3 and H3K27ac marks, respectively (Supplemental Fig. S6), with some variation across species owing to differences in data quality and alignability. The substantially greater abundance of H3K27ac elements was expected because the H3K27ac mark is associated with both active promoters and active enhancers, whereas the H3K4me3 mark is more specific to promoters. The two types of elements also had highly distinct distributions of state assignments, with the fully conserved state being the most frequent for the H3K4me3 mark but ranking much lower for the H3K27ac mark, beneath most single-species states (Supplemental Fig. S7). This difference suggests substantially lower rates of turnover in enhancers than promoters (see below).

### phyloGLM: Modeling of CRE turnover conditional on local genomic features

We addressed the third stage in our pipeline—modeling of gain/loss dynamics conditional on genomic features such as nearby gene expression or sequence conservation—with a new program, called phyloGLM, that allows the free parameters of the our phylogenetic model (*π* and *γ*) to be determined by a function of genomic features through a generalized linear model (GLM; Fig. 3). As shown below, this GLM-based design provides a rigorous framework for measuring the strength of association of individual genomic features with turnover rate, and for testing for differences in turnover rate between distinct groups of CREs (see Discussion).

**Figure 3:**
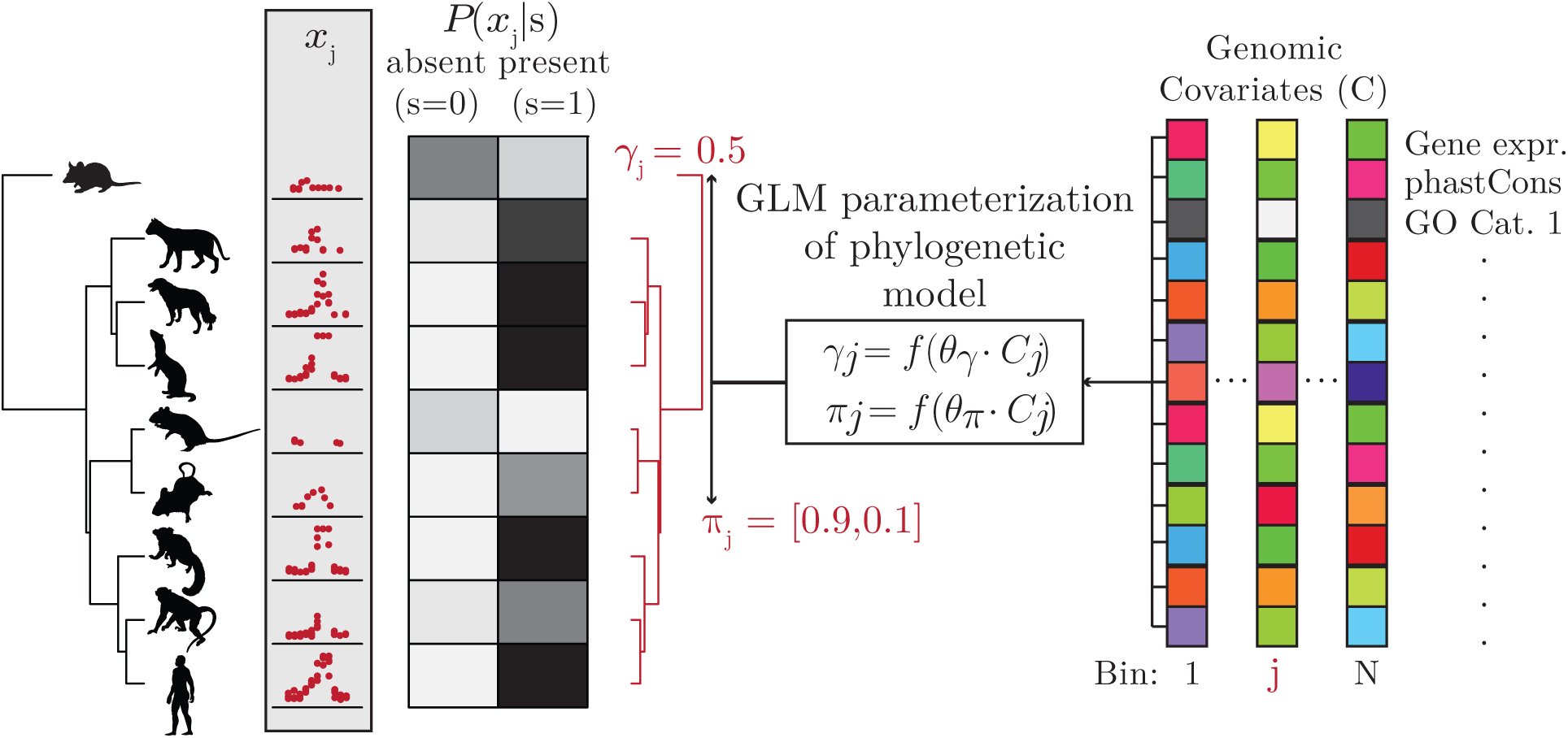
The phyloGLM model. phyloGLM combines the general modeling framework of Fig. 1 with a generalized linear model (GLM) to account for the influence of local genomic covariates (such as gene expression levels or sequence conservation scores) on the turnover process. In each genomic bin *j*, the turnover rate γ*_j_* and the equilibrium frequency for active elements π*_j_* are determined by a logistic function *f* (see **Methods**) applied to a linear combination of covariate values *C_j_* with weight vectors θ_γ_ and θ_π_, respectively. The phylogeny and hypothetical read counts *x _j_* for bin *j* are shown at the left, and hypothetical covariate values are represented by colored squares to the right. The shades of gray to the right of the phylogeny illustrates relative values of the conditional likelihoods *P*(*x_j_*|*s*).

### Application of phyloGLM to real data

We applied phyloGLM to the genome-wide predictions from epiPhyloHMM, separately analyzing the H3K4me3 promoter data set and two subsets of the H3K27ac data that correspond to likely promoters and likely enhancers. Thus, we were able to compare the enhancer data set with two distinct promoter data sets, one of which (H3K27ac) included more abundant but less precise predictions than the other (H3K4me3). To set up the analysis, we first assigned each CRE a putative target gene from Ensembl^52^ using simple distance-based rules, which essentially associated each CRE with the closest transcription start site (TSS) of a gene but discarded elements that could plausibly be associated with more than one gene (see **Methods** and Supplemental Fig. S8). Also based on proximity to the nearest TSS, we classified H3K27ac CREs as likely promoter (within 1.5 kb) or enhancer (within 100 kb) elements, and we similarly classified H3K4me elements as likely promoters (within 1.5kb) or discarded them. Finally, we associated each CRE with a collection of genic and cis-regulatory features that could potentially impact its turnover rate (Fig. S9). Broadly, these features described the numbers of CREs associated with the target gene, the expression patterns and annotated function of that gene, and measures of evolutionary constraint on the local DNA sequence. After removing all elements with incomplete covariate data, we were left with 5,368 H3K4me3 promoters, 7,220 H3K27ac promoters, and 25,673 H3K27ac enhancers for further analysis. These features were somewhat correlated with one another, but most correlations were weak (Supplemental Fig. S10). We fitted phyloGLM separately to these three sets of elements, estimating all free parameters by maximum likelihood and conditioning on a phylogeny with branch lengths based on published estimates of divergence times in millions of years^53^. In all cases, we separately parameterized the branch to the outgroup (opossum), on which gains and losses are difficult to distinguish, to avoid skewing the other parameter estimates.

This model permitted us to compute the expected total numbers of gains and losses along each branch of the phylogeny, conditional on the data and the fitted model. Similar patterns of gain and loss were observed for the H3K4me3 and H3K27ac promoters, with fewer total events in the H3K4me3 elements (0.97 vs. 1.22 events per element on average; Supplemental Fig. S11). For H3K27ac elements, we found that the overall rates of gains and losses are fairly similar, with somewhat more gains than losses, both within promoters and within enhancers (Supplemental Fig. S12). However, the total rate of turnover for enhancers appears to be approximately twice that of promoters, with 2.47 events per element compared with 1.22 for the H3K27ac elements. The numbers of expected events per branch were roughly proportional to the branch lengths, with long branches tending to be assigned more events than short branches. The gain/loss proportions were somewhat variable across branches, but this variation likely reflects a combination of true differences and biases from human-referenced alignments and differences in ChIP-seq data abundance and quality across species (see Discussion).

A comparison of the distributions of numbers of events per CRE provided further support for a roughly two-fold higher rate of turnover at enhancers than promoters, with median values of 0.0075 and 0.0031 events per million years (myr), respectively, for the H3K27ac data (Fig. 4A; *p* < 2.2 × 10^−16^; Wilcoxon signed-rank test). From these distributions and the estimated phylogeny, it was also possible to estimate a distribution of the “half-life” (time required for half of active elements to be lost) for each type of CRE. For the H3K27ac data, the median half-life for enhancers is 130 myr and that for promoters is 552 myr (Fig. 4B; see **Methods**). These estimates are substantially lower than previous estimates of 296 myr and 939 myr, respectively^35^. However, our estimate of the median turnover rate for promoters based on the less noisy H3K4me3 data set was ∼30% lower, at 0.0022 events per myr, corresponding to a half-life estimate of 937 myr, in much better agreement with the corresponding previous estimate. Thus, it seems likely that our H3K27ac turnover rate estimates are substantially inflated by the lower resolution ChIP-seq data (see Discussion).

**Figure 4:**
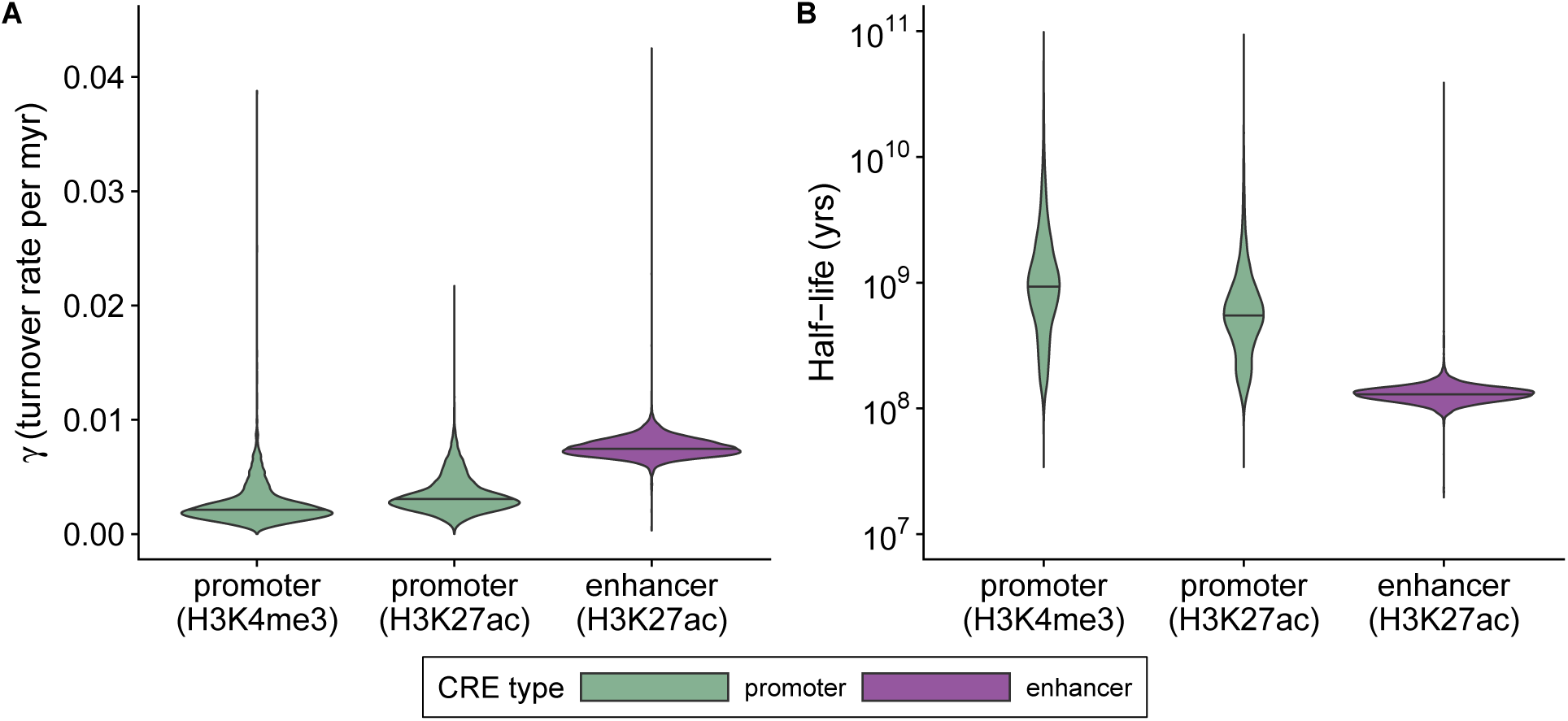
Enhancers show greater rates of turnover than promoters. (A) Distributions of estimated turnover rates per bin (γ*_j_*) across all elements, based on the full phyloGLM model. Results are shown for separate analyses of promoters based on the H3K4me3 mark, promoters based on the H3K27ac mark, and enhancers based on the H3K27ac mark (see **Methods**). The phylogeny and branch lengths (in millions of years) were obtained from ref. ^53^. The branch to the opossum outgroup was excluded in rate estimation. To simplify the visualization, a small number of elements (41 of 38,170) with turnover rates > 0.044 were omitted. The bar in each violin plot represents the median. (B) Similar distributions of times for half of all active elements to decay to an inactive state (half-life).

### Gene-expression-related features associated with turnover rates

In addition to allowing us to characterize overall rates of turnover, our GLM-based framework enabled us to examine the strength and directionality of the association between each of the genomic features we considered and the rates of turnover at enhancers and promoters. We begin by considering these relationships for gene-expression-related features, and examine the remaining features in subsequent sections. For promoters, three gene-expression-related covariates had statistically significant associations with turnover rate: the level of expression of the target gene in the liver (the assayed tissue here), the number of tissues in which the target gene was expressed, and the cross-tissue expression dispersion (Fig. 5). For liver expression and the number of tissues, increased values of the covariate were associated with significantly decreased turnover rates. These observations are broadly consistent with a variety of previous analyses that have indicated that CREs associated with high levels of expression in the tissue of interest or with broad expression patterns across tissues tend to experience elevated levels of constraint^8, 35, 54–56^. The positive correlation with cross-tissue expression dispersion (significant for H3K4me3 only) appears to reflect a similar trend.

**Figure 5:**
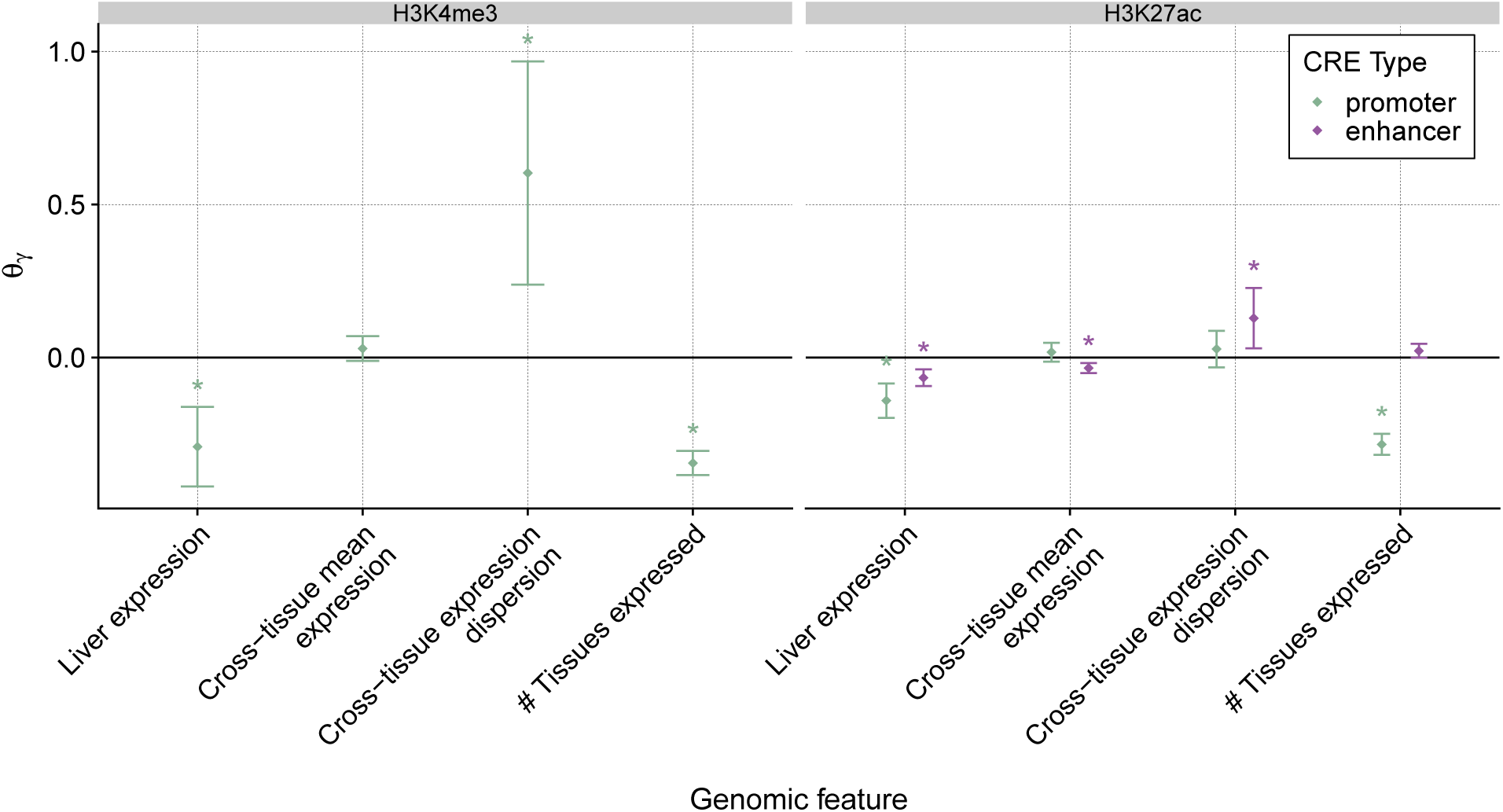
Marginal effects of gene-expression-related features on turnover rates. Estimates of the coefficients in the generalized linear model (θ_γ_) for four gene-expression-related features, for promoters based on H3K4me3 (left) and both promoters and enhancers based on H3K27ac (right). A positive value indicates that a feature increases the turnover rate and a negative value indicates that it decreases the turnover rate. The branch to the outgroup is excluded from rate estimation. Error bars represent approximate 95% confidence intervals (see **Methods**). “*” indicates a significant difference from zero (*p* < 0.05 after Benjamini-Hochberg correction) based on a likelihood ratio test.

The results for enhancers were generally similar to those for promoters, but tended to be somewhat weaker, likely in part owing to the difficulty of correctly linking enhancers with target genes. One notable difference for enhancers was that the number of tissues in which the target gene was expressed was not significantly associated with turnover rate. This difference might result in part from tissue-general “housekeeping” tending to have fewer enhancers than tissue-specific genes^57^. In addition, housekeeping genes are likely enriched for proximal enhancers, which tend to be excluded by our filters.

### Additional features associated with turnover rates

The remaining genomic features describe either measures of sequence constraint of the CRE (phastCons^6^) or the gene (pLI^58^), or aspects of the local regulatory “architecture” of each target gene, including the numbers of enhancers and promoters and the distance of each enhancer from the TSS (Fig. 6). Both enhancers and promoters displayed a negative correlation between CRE sequence conservation, as measured by phastCons, and turnover rate. As previously noted^24, 35, 56^, this observation indicates that elements that are more constrained at the DNA sequence level are also more resistant to evolutionary gain and loss. Interestingly, however, we observed no significant correlation between constraint against loss-of-function variants in the gene, as measured by pLI scores, and turnover rates of associated enhancers or promoters (see next section).

**Figure 6:**
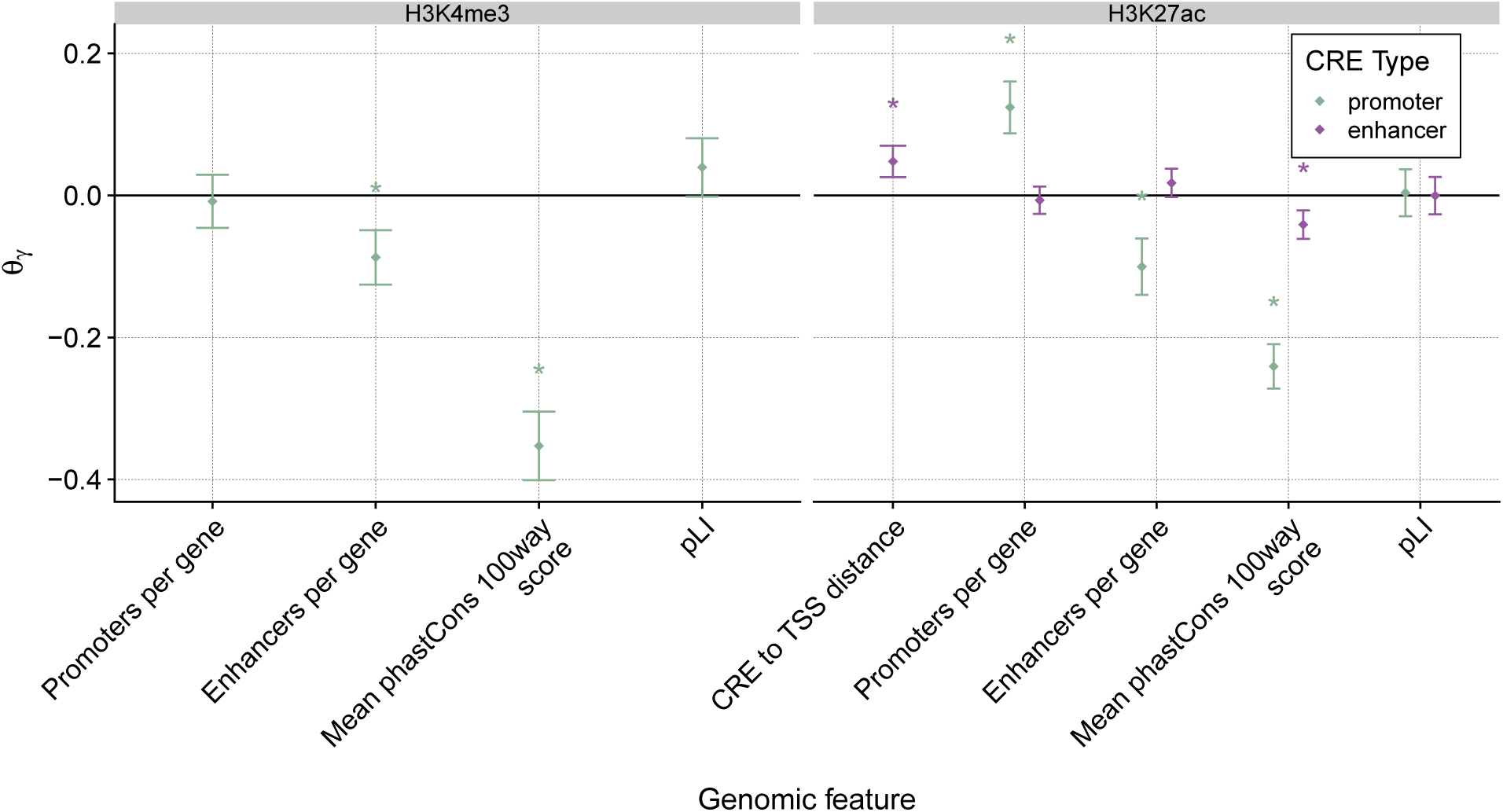
Marginal effects of remaining features on turnover rates. Estimates of the coefficients in the generalized linear model (θ_γ_) for four additional features, for promoters based on H3K4me3 (left) and both promoters and enhancers based on H3K27ac (right). A positive value indicates that a feature increases the turnover rate and a negative value indicates that it decreases the turnover rate. The branch to the outgroup is excluded from rate estimation. Error bars represent approximate 95% confidence intervals (see **Methods**). “*” indicates a significant difference from zero (*p* < 0.05 after Benjamini-Hochberg correction) based on a likelihood ratio test.

Among the architectural features, the strongest correlate at the enhancer level is the distance to the TSS, a quantity that is positively associated with turnover rate. As has been noted in several recent studies^24, 35, 56^, this increased constraint against turnover on enhancers close to the TSS likely reflects an enrichment for genuine enhancer-gene interactions and direct influence on the expression of the target gene. Another observation that echoes a previous finding is that the number of enhancers per gene is positively correlated with the enhancer turnover rate but negatively correlated with the promoter turnover rate. As previously noted^24^, this observation suggests that larger ensembles of enhancers associated with the same target gene tend to impose additional constraints against promoter turnover, but nevertheless to relax constraint on each of the enhancers themselves, perhaps because each enhancer is less essential to the overall regulatory architecture of the locus (see also refs. ^59–61)^. We also observed significant effects for several top-level Reactome categories, suggesting that biological function may provide additional information about regulatory constraint (Supplementary Fig. S13). Together, these observations suggest that CREs evolve in a manner that is strongly dependent on the local regulatory context in which they appear.

### Differences between DNA sequence and epigenetic conservation in CREs

The correlation of turnover rate with the phastCons scores of CREs but not with the pLI scores of target genes is curious, but what does it signify? The pLI score for a gene measures the probability of intolerance to (generally heterozygous) loss-of-function mutations in that gene, as inferred from patterns of variation in ultra-deep human exome sequencing data^58^. pLI scores can be used to differentiate between haploinsufficient and haplosufficient genes or, similarly, between dosage-sensitive and insensitive genes^58^ (although, strictly speaking, the scores are directly informative only about the strength of selection acting on heterozygotes^62^). By contrast, phastCons scores simply measure a reduction in fixed derived alleles, and do not effectively differentiate among various forms of negative selection. Therefore, the observed difference in correlation suggests that gains and losses of CREs are generally deleterious but perhaps do not depend strongly on the dominance or dosage properties of target genes.

To investigate this issue further, we examined CRE turnover rates at the promoters of two classes of genes that serve as proxies for dosage insensitivity and sensitivity, respectively: genes that encode proteins involved in metabolism such as enzymes (whose action tends to be relatively insensitive to protein abundance) and genes that encode proteins that regulate gene expression such as transcription factors (which tend to be more sensitive to abundance)^63, 64^. We found no significant difference between these classes of genes in the turnover rates of CREs in either promoters or enhancers (Fig. 7A), further supporting the idea that turnover rates have little dependency on dosage or dominance. Interestingly, however, when we condition on the expected number of turnover events at each CRE and focus on the CREs that have undergone the fewest events, we do observe increased sequence conservation at the more dosage-sensitive regulatory genes. This difference is observed both for promoters (Fig. 7B) and enhancers (Fig. 7C), although it is statistically significant for promoters only. It suggests that, while both classes of genes are similarly resistant to mutations that result in the complete gain and loss of elements (Fig. 7A), dosage-insensitive genes are more tolerant of nucleotide substitutions that do not result in complete gain or less events (Fig. 7B&C). These observations illustrate how the evolutionary dynamics of CRE gain and loss may differ from those for nucleotide substitutions owing to the pronounced effects of turnover events on gene expression, and, more generally, how patterns of evolutionary constraint across the genome may depend on the effect sizes of mutations.

**Figure 7:**
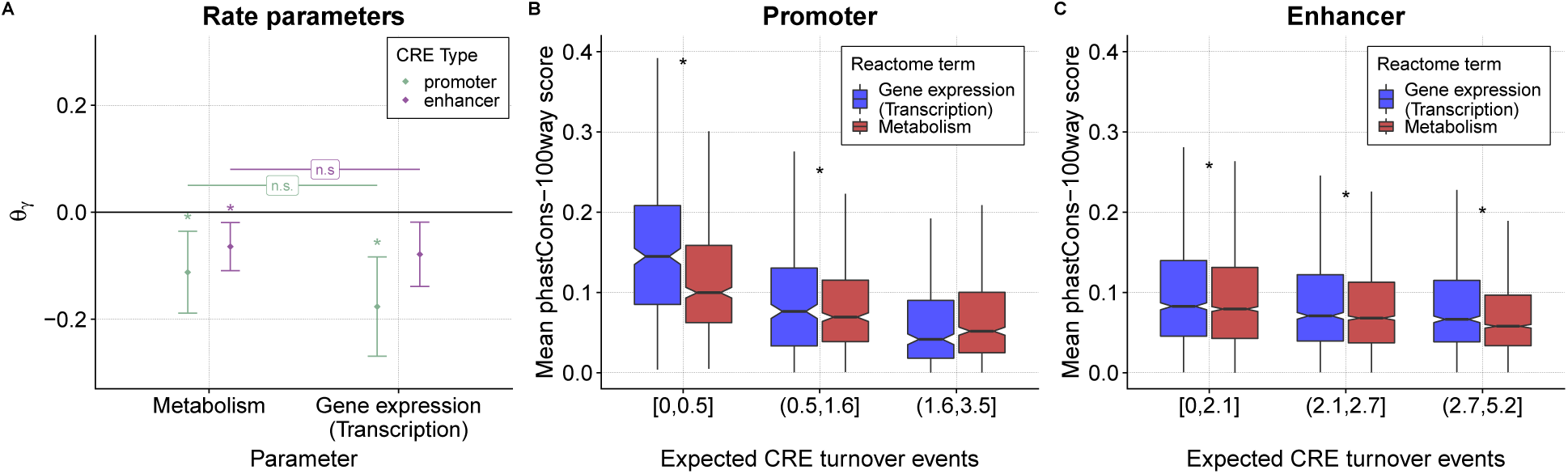
Turnover rates and sequence conservation at CREs for dosage-sensitive and insensitive genes. (A) Estimated coefficients *θ_γ_* for indicators for the Reactome^94^ categories “Metabolism” (less dosage-sensitive) and “Gene expression (Transcription)” (more dosage-sensitive) at associated promoters and enhancers (both based on H3K27ac). Error bars represent approximate 95% confidence intervals (see **Methods**). “*” indicates a statistically significant difference from zero (*p* < 0.05 after Benjamini-Hochberg correction) based on a likelihood ratio test. The differences between the two categories of genes are not significant (n.s.). (B) Mean phastCons scores for vertebrates (100-way alignment in UCSC Genome Browser) in H3K27ac-based promoters of the same two categories of genes, stratified by the expected number of turnover events per CRE. Notches correspond to 1.58 · *IQR*/ *√n* indicating an approximate 95% confidence interval for the median. “*” indicates a significant difference between gene categories (*p* < 0.05, *t*-test with Benjamini-Hochberg correction). (C) Same as *(B)* but for H3K27ac-based enhancers. For visual clarity, outliers (more than 1.5 · *IQR* from the hinge) are not shown in (B) and (C).

## Discussion

In this article, we have introduced a new probabilistic modeling framework for inferring the dynamics of CRE gain and loss, which accounts for phylogenetic correlations among species, uncertainty in peak calls from ChIP-seq data, and the influence of local genomic features on turnover rates. By applying our methods to H3K4me3 and H3K27ac ChIP-seq data^35^, we find support for a number of previously reported results, including a substantially higher rate of turnover in enhancers than promoters, negative correlations of turnover rate with DNA sequence conservation, expression level, and tissue breadth, positive correlations with distance from the TSS, and a strong dependency on features of the local regulatory architecture such as number of enhancers per gene. Overall, we find that enhancers are gained and lost at about twice the rate of promoters during mammalian evolution, with median rates of 0.0075 and 0.0031 events per element per million years, respectively, based on the H3K27ac data.

In addition, we made use of our modeling framework to examine an apparent lack of correlation between turnover rates at CREs and the haploinsufficiency or dosage sensitivity of target genes, as measured approximately using pLI scores^58^. We found that turnover rates were significantly negatively correlated with DNA sequence conservation in CREs, suggesting that both whole gain/loss events and nucleotide substitutions are deleterious, but that turnover rates were not correlated with pLI scores, suggesting that gain/loss events are no more deleterious at haploinsufficient/dosage-sensitive genes than at haplosufficient/dosage-insensitive genes. However, when we conditioned on the expected number of turnover events at each CRE, a positive correlation became evident between sequence conservation and dosage sensitivity at low-turnover CREs (Fig. 7). We interpret this result as indicating that DNA substitutions that do not cause complete gain or loss events are more easily tolerated at dosage-insensitive genes than at dosage-sensitive genes. These mutations of small effect at dosage-insensitive genes may be allowed to accumulate and perhaps compensate for one another, permitting drift in CRE sequences as long as it does not cause the gain or loss of a whole element. By contrast, gains and losses of entire CREs have sufficiently large effects that they are deleterious at both dosage-sensitive and -insensitive genes. Thus the divergence between epigenetic and sequence constraint is potentially informative about the mode of selection at each locus. These observations may help to explain previous reports of CREs that display conservation of epigenetic marks but not the DNA sequence^65–68^ (see also refs. ^69, 70^).

For various reasons, we have approached the problems of identifying regulatory elements and modeling their evolution separately, using the epiPhyloHMM and phyloGLM programs, respectively. This strategy allows us to address the problem of segmenting the genome into “active” and “inactive” regions in a relatively efficient manner, using a simpler model, and then characterize the turnover process using a richer model that conditions on a diverse collection of genomic features. It also has practical advantages in terms of modularity of software development and efficient processing of genome-wide data. At the same time, this strategy has the limitation that the richer evolutionary model implemented in phyloGLM is not exploited in element identification, which in principle, could result in loss of power. Still, this limitation does not appear to be of major practical importance because identifying regions containing active elements turns out to be fairly straightforward. A related limitation is that we analyze the genome in bins of fixed size, which occasionally results in spurious inferences of turnover when the boundaries of the bins align poorly with the locations of peaks. Nevertheless, this simple approach is generally fairly effective for pooling read counts along the genome and accommodating limitations in the genomic resolution of peak locations (see below).

Notably, our flexible phyloGLM design allows not only for improved modeling of evolutionary dynamics, but also direct assessment of hypotheses about how these dynamics depend on various aspects of genomic context, such as gene expression, local regulatory architecture, and sequence conservation. Since phyloGLM considers all covariates together in a single model, we avoid the need for complex post-hoc analyses, for example, that require matching of foreground and background sets of elements in terms of relevant covariates. Similar approaches have been used for the estimation of *dN*/*dS* rates^71^ and probabilities of fitness consequences for new mutations^55, 72^, but to our knowledge, this approach has not been previously employed in the study of CRE evolution.

Two other recently published methods^40, 42^ have addressed the problem of inferring evolutionary dynamics from multi-species epigenomic data using strategies that are similar to ours, but are also different in key respects. Importantly, both of these methods avoid separating the element identification and evolutionary inference problems—a decision that has potential advantages, provided the data have sufficiently high resolution to avoid overfitting, but that is also costly in computational efficiency. The first method^42^ directly models an evolving continuous signal (replication timing, in their application) along a collection of aligned genomes using an elegant combination of a branching Orstein-Uhlenbeck process along a phylogeny and a hidden Markov model along the genome sequence, which is fitted to genome-wide data by expectation maximization. This approach appears to be quite powerful but it differs from ours in that it focuses on direct modeling of a continuous molecular phenotype, rather than on describing a relationship between a discrete property of interest (such as transcription factor binding or CRE activity) and functional genomic data describing that property (such as ChIP-seq read counts). The strategy has potential advantages when the property of interest truly is continuous, but could also tend to overfit a complex signal. The second method^40^ is more similar to ours in that it does distinguish between discrete “positive” and “negative” states (in this case, reflecting DNA methylation status), again using a combination of a hidden Markov model and a phylogenetic model. The structure of this HMM is more complex than ours, however, and requires Monte Carlo methods for fitting. The setting for this method is also different from ours in that the whole genome bisulfite sequencing data being analyzed appear to provide a higher-resolution, more precise readout of the feature of interest, avoiding some of the uncertainty of peak-calling from ChIP-seq data. More generally, modeling of comparative epigenomic data remains an active area of research, with a number of newly developed methods that make similar but complementary modeling assumptions, and more work will be needed to find which approaches are best suited for various applications of interest.

A major problem faced by all current modeling approaches—and a reason why model-free methods have often been used instead—is the fundamental imprecision of comparative epigenomic data. In most data sets, there is considerable uncertainty not only in the strength of the signal along the genome, but also in the precise genomic position and breadth of that signal (i.e., in peak height, location, and width, in the context of ChIP-seq data). This uncertainty is compounded by errors in alignment and orthology identification between species. Evolutionary models must therefore strike a balance between getting the most out of the available data, and avoiding biases that come from assuming unrealistic levels of precision or resolution. Indeed, in practice, all of the current modeling approaches have required the use of extensive heuristics and filters before and/or after they are applied. In our case, the imprecision of the data resulted in a tendency to fragment individual elements into multiple predicted segments, for example, because peaks did not align well in width or position across species. We attempted to mitigate this problem by postprocessing the epiPhyloHMM predictions with heuristic rules that joined and filtered elements (see **Methods**). Nevertheless, these misalignment and fragmentation issues undoubtedly produced some upward bias in our estimates of turnover rate. This effect most likely drives the substantial elevation in rate for the promoters based on the noisier H3K27ac data as compared with the more precise H3K4me3 data. More work will be needed both to improve the precision of comparative epigenomic data and to accommodate uncertainty in models of evolutionary dynamics.

Two other limitations of our analysis that are broadly shared with other comparative genomics studies concern the use of reference-based multiple alignments and proximity-based rules for associating CREs with target genes. Our use of human-referenced multiple alignments (which are also used in refs. ^40^ & ^42^) prevents us from analyzing portions of the other genomes that do not align to the human genome, and therefore creates a general bias toward “gains” in human and closely related species, and toward “losses” in more distant parts of the tree, as is evident in a close inspection of Supplementary Figs. S11 & S12. This same limitation makes it infeasible to test for differences in gain or loss rates across branches or clades of the phylogeny, because alignment-induced biases would likely overwhelm real biological differences. There has been some progress in recent years toward generalized reference-free multiple alignment methods^73, 74^ but much more work is needed on this important problem.

Similarly, the linking of CREs, particularly enhancers, with target genes is another fundamental unsolved problem that pervades many genomic analyses. In our case, it is likely that a substantial fraction of enhancers are mis-assigned a target gene, with downstream effects on a number of our analyses (e.g., Figs. 5, 6, & 7). Experimental work to link enhancers to the correct target genes, either via 3D-chromatin capture^75–77^, or large scale genome editing^78^, will help to improve this issue over time.

## Acknowledgments

This research was supported by US National Institutes of Health grants R01-HG010346 and R35-GM127070. The content is solely the responsibility of the authors and does not necessarily represent the official views of the US National Institutes of Health.

## Software availability

epiPhylo is available as an R package at https://github.com/ndukler/flexPhyloHMM. phy-loGLM is also available as an R package at https://github.com/ndukler/phyloGLM.

## Methods

### ChIP-seq data preparation

All ChIP-seq data were obtained from ref. ^35^. Reads were aligned to the reference genome for each species^79–86^ (obtained from the UCSC genome browser) using bowtie2 (v 2.2.9)^87^. Each read was summarized by the single base at the center of the read and converted to the human reference genome using the liftOver utility and the best reciprocal chains supplied by the UCSC genome browser^88^ (hg38). A coverage map was computed from the converted reads and summed to get the total number of reads per 250bp bin. Regions that could be aligned to the human genome from one or more other species were combined if they were less than 50kb apart and were expanded by 5kb to either side to create genomic blocks within which to run the phylo-HMM. Sections of the human genome that were not covered by one of these blocks were excluded from the analysis, leaving 2.83Gb for further analysis.

### Model for peak-calling

Our peak-calling model has two versions: the full model used in epiPhyloHMM and a simpler two-state model that is applied separately to the data for each species in a preprocessing step (as detailed below). The two versions are the same except for the state space. In the two-state model, the probability of the data in bin *j*, for presence/absence state *p* ∈ {0, 1} is given by a negative binomial mixture model:

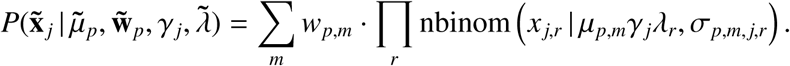

Here, x̃*_j_* is a vector of the read counts for each replicate *r*, *w_p_,_m_*is the weight of mixture component *m* for state *p* (such that ∑*_m_ w_p_*_,*m*_ = 1), µ*_p_*_,*m*_ is the mean count for state *p* and component *m*, σ*_p_*_,*m*,_ *_j_*_,*r*_ represents the dispersion parameter for the negative binomial distribution (see below), γ*_j_* is the fraction of bases in bin *j* that are aligned to the human reference genome, and λ̃ = {λ*_r_*} is a set of scaling factors that account for the sequencing depth of each replicate *r*. As noted in the Results section, we use a three-component mixture model for the “presence” state (*p* = 1) and a single component for the “absence” state (*p* = 0). The scaling factor λ*_r_* is calculated as:

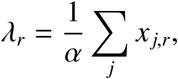

where α = max*_r_* (∑*_j_ x_j,r_*). To account for differences in the dispersion of read counts for different mean depths, we make use of the model from DESeq2 (ref. ^47^) where the dispersion of the distribution σ is defined as a function of the mean µ and two free parameters, θ_1_ and θ_2_:

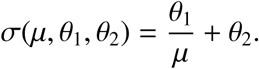

We use the DESeq2 software to estimate θ_1_ and θ_2_, after sub-sampling the genome to obtain roughly similar numbers of low-, medium-, and high-coverage sites. We then calculate the dispersion per data-point as,

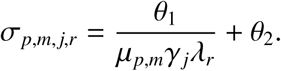

This strategy accounts for effects of sample library depth and cross-species alignability on the expected read-count depth for each state *p*, mixture component *m*, replicate *r*, and bin *j*.

The transition model is a simple two-state model with auto-correlation parameters ρ_1_ for the peak state and ρ_0_ for the background state (Fig. 2).

### Hidden Markov model

The hidden Markov model used by epiPhyloHMM includes a set of states {*s*_1_, …, *s_E_*} representing all patterns of CRE presence and absence at the tips of the tree, up to a maximum number of gain/loss events (three, in our application). Conditional on the state (and, implicitly, on the gain/loss history), the read-counts at the tips of the tree are independent. Thus, the emission probability for the observed data in state *s_e_* at site *j* is given by a product over tips (species) *t* and presence/absence states *p*:

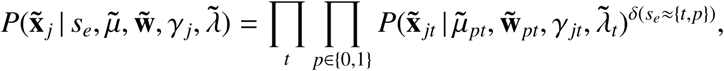

where δ(*s_e_* ≈ {*t*, *p*}) takes a value of one if HMM state *s_e_* is consistent with species *t* having presence/absence state *p* and a value of zero otherwise. Where there is a large alignment gap in a species (with size ≥5 kb), we force the emission probability for “presence” (*p* = 1) in that species to zero, presuming a deletion.

The matrix of transition probabilities consists of three different types of transitions. First, self-transitions for all active and inactive states, have probabilities ρ_1_ and ρ_0_, respectively (Fig. 2). Second, all transitions from active states to the inactive state have probability 1 − ρ_1_. Third, each transition from the inactive state to any active state *s_e_* has probability 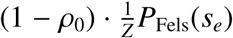. Here, *P*_Fels_(*s_e_*) is the probability of the presence/absence pattern at the tips of the tree consistent with state *s_e_*, as computed using Felsenstein’s pruning algorithm, under our phylogenetic turnover model (implicitly conditioned on the given phylogeny and the parameters π and γ). Because not all presence/absence patterns are possible, this probability must explicitly be normalized by the sum across all allowable states, *Z* = ∑_active *s*_*_e_ P*_Fels_(*s_e_*). Thus, the relative frequencies of the active states are proportional to their equilibrium probabilities under the specified phylogenetic process. All other elements of the transition matrix are fixed at zero, preventing direct transitions between active states.

### epiPhylo model fitting

For reasons of efficiency, we fit the epiPhyloHMM model to the data approximately in several successive steps. We first fit the peak-calling models separately to the data for each species, using a subset (125 Mb) of the mapped reads and estimating all free parameters by maximum likelihood with the L-BFGS-B algorithm^89^. As noted above, this calculation made use of the dispersion model that was pre-estimated using DESeq2. We then split the multi-species data into twenty partitions of similar size, with break-points in 50kb-long regions lacking alignment to the human genome by any other species. Separately in each species, we converted bins with <15% of bases aligning to the human genome to missing data. We then estimated the ρ_1_, ρ_0_, π, and γ parameters of the epiPhyloHMM model by maximum likelihood using the LBFGS-B algorithm^89^, keeping the species-specific peak-calling parameters—namely, the mixture coefficients *w* and mean counts µ⃗—at their previously estimated values. We then obtained an inital set of element calls using the Viterbi algorithm, and filtered them by the following heuristics:

1. We grouped maximal sets of elements that were separated by at most one “background” bin (as sometimes occurs due to the sparse design of the transition matrix; Fig. 2).
2. For each grouped element, we computed an alignment “score” equal to the sum of alignment scaling factors γ*_j_* across all bins *j* in the element and across all species.
3. We retained the element in the group having the highest alignment score.
4. In addition, we retained any elements having a score that exceeded a designated threshold *T* (*T* = 16 in our analysis).
5. We masked any remaining elements from the data by re-setting their alignment scale factors γ*_j_* to 0.

After this masking step, we re-estimated the ρ_1_, ρ_0_, π, and γ parameters of the epiPhyloHMM model separately for each partition of the data. Then we obtained our final set of predictions by running the Viterbi algorithm genome-wide using the median values of these per-block estimates.

### CRE-gene association and annotation as enhancers and promoters

All assignments were based on distances to genes obtained from Ensembl build 93 (ref. ^52^) via BiomaRt^90, 91^. Promoter regions were defined per transcript as the interval ±1.5kb of the annotated transcription start site (TSS). Each promoter region was associated with the gene linked to the TSS in question, but multiple promoter regions were allowed per gene. H3K4me3 elements that overlapped (by at least one nucleotide) with a single promoter region were annotated as ‘promoter’ and associated with the corresponding gene. H3K4me3 elements that overlapped with multiple promoter regions were annotated as an ‘unassociated promoter’ (promoter UA). H3K4me elements that did not overlap with any promoter regions were annotated as ‘unknown’ (unk). For H3K27ac marks, the same rules were used to label an elements as promoter or promoter UA.

To classify enhancers, we first defined ‘expanded promoter regions’ as intervals ±10kb of the TSS, again merging them across transcripts of the same gene. H3K27ac elements that overlapped with a single expanded promoter region were annotated as a ‘proximal enhancer’ (enhancer proximal) and associated with the corresponding gene. H3K27ac elements that overlapped with multiple expanded promoter regions were annotated as an ‘unassociated proximal enhancer’ (enhancer proximal UA). H3K27ac elements that did not overlap with any expanded promoter regions but still fell within 100 kb of a TSS were annotated as a ‘distal enhancer’ and associated with the closest gene (enhancer distal). H3K27ac elements that met none of these criteria were labeled as ‘unknown’. This scheme is represented in Supplemental Fig. S8.

### Linking phylogenetic parameters to genomic features for phyloGLM

As described in the **Results** section, the phylogenetic model for CRE turnover is defined by two free parameters: the turnover rate, γ, and the equilibrium frequency of element “presence” distribution, π. These parameters are defined as generalized linear functions of a vector of genomic features, or covariates, that are assumed to be available for each bin *j*. Specifically, the turnover rate, γ*_j_*, for a given bin *j* with covariate vector *C_j_* is defined by passing a linear combination of *C_j_* and a vector of coefficients θ_γ_ through the logistic function,

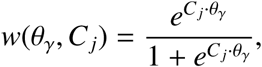

with bounds of γ_min_ ≤ γ*_j_* ≤ γ_max_ imposed as follows,

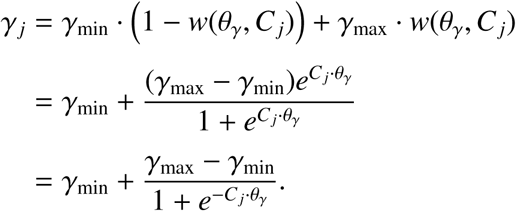

Our implementation allows for a separate set of coefficients θ_γ,*e*_ for each edge *e* of the phylogeny, but in practice we separate only the branch to the outgroup and apply the same coefficients to all other branches of the tree. This strategy ensures that weak power to distinguish gains and losses on this branch, and poor alignability to the outgroup, do not drive the maximum likelihood estimates for other parts of the tree.

Similarly, the stationary distribution of element presence π*_j_* is defined as

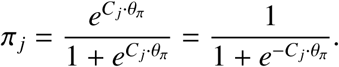

In order to prevent numerical underflow, if π*_j_* < 10^−3^, we reset π*_j_* = 10^−3^; similarly, if 1−π*_j_* < 10^−3^, we reset π*_j_* = 1 − 10^−3^.

### Fitting a phyloGLM model

The phyloGLM model was fit to the data using the L-BFGS-B algorithm^89^ as implemented in the ‘optim’ function in R^92^. Gradients with respect to the log likelihood ℒ(θ⃗; **x⃗**) (where θ⃗ denotes the entire parameter set) for elements of the coefficient vectors θ_γ_ and θ_π_ were computed using the chain rule. For example, the gradient for an element *i* of the vector θ_γ_, denoted θ_γ,*i*_, is given by,

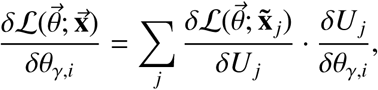

where the sum is across all bins, ℒ(θ⃗; **x⃗**_j_) represents the contribution of bin *j* to the log likelihood function, and *U_j_* = θ_γ_ · *C_j_* is the linear combination of features for bin *j*. The partial derivative 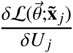 was computed numerically. However, the final term can easily be computed analytically as,

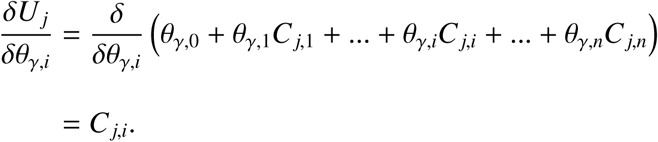

Notice that the use of the chain rule considerably accelerates these calculations because 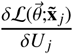 only needs to be computed once per phylogenetic parameter (γ and π), and then can be propagated efficiently to all of the individual coefficients. The same approach works for elements of θ_π_.

### Preparation of genomic features

Distances to the TSS were based on annotations from Ensembl build 93 (ref. ^52^), accessed via BiomaRt^90, 91^. The distance was computed from the nearest boundary of the CRE to the annotated TSS position. Gene expression features were based on data downloaded from the GTEx web portal (https://gtexportal.org/). Mean phastCons-100way scores were calculated using the GenomicScores package^93^. pLI scores were collected from ftp://ftp.broadinstitute.org/pub/ExAC_release/release1/manuscript_data/. Functional annotations of genes were obtained from Reactome 2018^94^. For the fitting of the phyloGLM model, only bins that were unambiguously associated with a single gene and had no missing covariate values were used. After this filtering, 7,220 promoters and 25,990 enhancers associated with 5,307 and 5,552 unique genes remained. All non-categorical CRE covariates were scaled to have mean 0 and standard deviation 1 to improve model-fitting performance and produce comparable coefficient values across covariates.

### Expected numbers of gains and losses per branch

We calculated the expected numbers of gains and losses per branch (Supplemental Figs. S11 & S12) using the standard message-passing algorithm on the phylogeny, followed by estimation of the probabilities of each state transition per branch (see ref. ^95^ for details).

### Half-life estimation

Under our model, the instantaneous rate of transition from the active state to the inactive state is given by γ(1 − π). Thus, the half-life 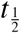, or time required for half of active elements to become inactive, is given by:

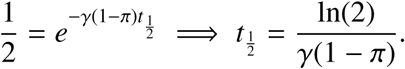

### Dosage-sensitivity analysis

Annotations of genes as “Metabolic” or “Generic Transcription Pathway” were obtained from Reactome 2018^94^. Only genes having one of the functional annotations “R-HSA-1430728” or “R-HSA-212436” were selected for analysis; genes with both labels were omitted. To test for significant differences between the two gene categories, we compared the log likelihoods of phyloGLM models having (i) separate coefficients for the two gene categories and (ii) one coefficient that applied to both categories, computing *p*-values for one degree of freedom. Mean 100way phastCons scores were calculated using bwtool^96^.

## Supplemental Materials

### Simulation study

To test the power of our model to recover the true peak calls and parameters, we simulated data under the model used for inference, allowing for all 2*^N^* states, given a variety of genome sizes, phylogenetic trees, and values of the turnover parameter (γ). We fixed the parameters for the peak calling model to their true values to mimic our multi-step fitting process and isolate the ability of the model to correctly estimate the evolutionary parameters of interest.

With regard to calling peaks at the species level, we found that epiPhylo performed well by both precision and recall metrics but suffered slightly at higher turnover rates (Fig. S1). Model predictions were evaluated on a per-bin basis so that a prediction was considered a true prediction if the model’s state call for a bin in a particular species matched that bin’s state in the simulation. Each bin in a multi-bin element was evaluated separately. Across a broad range of rate parameters and tree topologies, estimates of γ converged to an inflated, but stable estimate of γ with as little as 250 kb when γ was large and as much as 250 Mb when γ was small (Fig. S2). The stationary probability for an absent element (1 − π) was systematically underestimated while estimates of the autocorrelation ρ_1_, converged to the true values given as little as 2.5 Mb of data (Fig. S3-S4). Systematic biases in the fitted values of γ and π occur to accommodate due to a ridge in the likelihood surface (Fig. S5), however this does not greatly impact the recovery of true regulatory elements in our simulation.

## Supplemental figures

**Figure S1:**
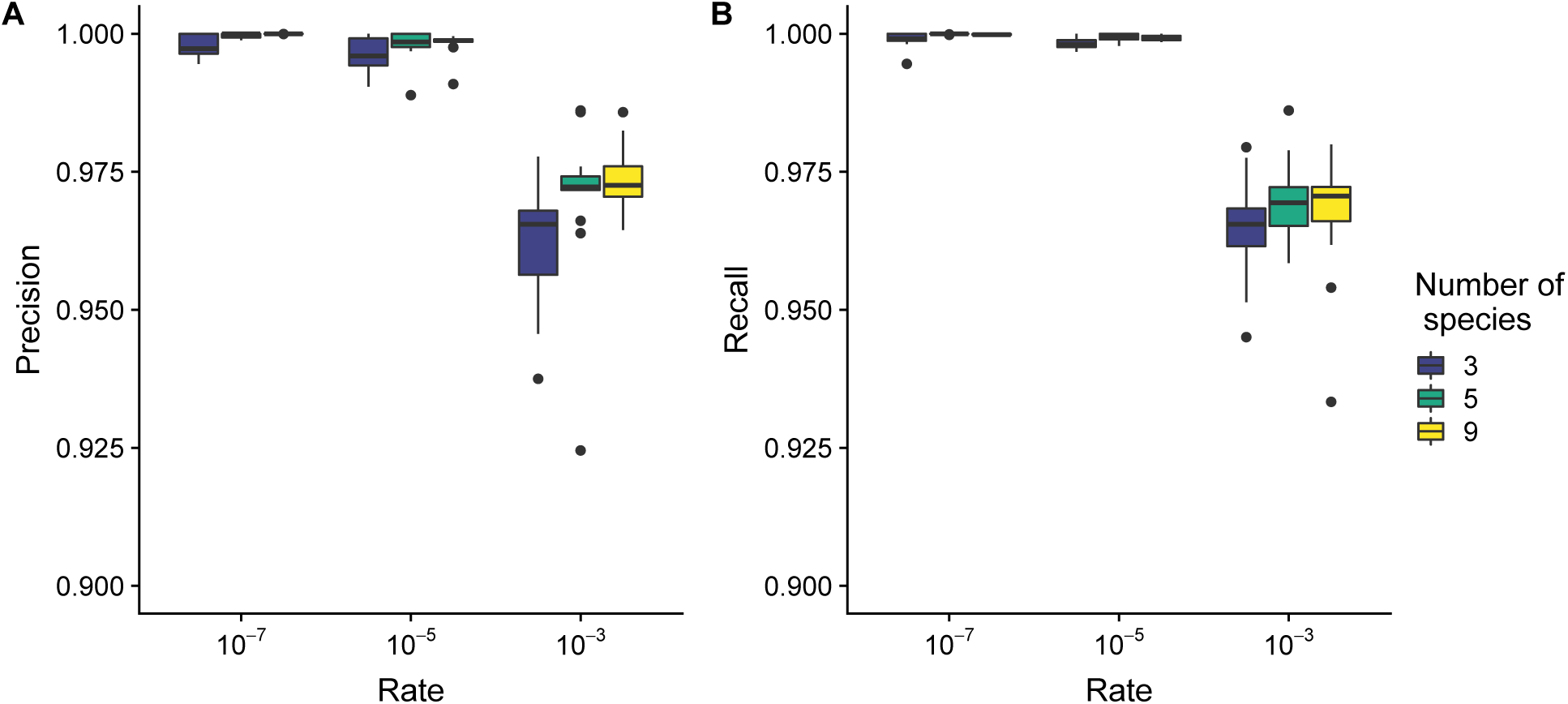
EpiPhylo performance as a function of rate for simulated data. (A) Precision of the epiPhylo model based on the number of 250bp bins for each species with correct state calls across three different trees. (B) Recall of the epiPhylo model based on the number of 250bp bins for each species with correct state calls across three different trees. For both calculations, the state that epiPhylo calls is used to assign active elements across species which are then compared to the true peaks at the species level. All possible species configurations were simulated, and the epiPhylo model was fit with all configurations that required three or fewer mutations.

**Figure S2:**
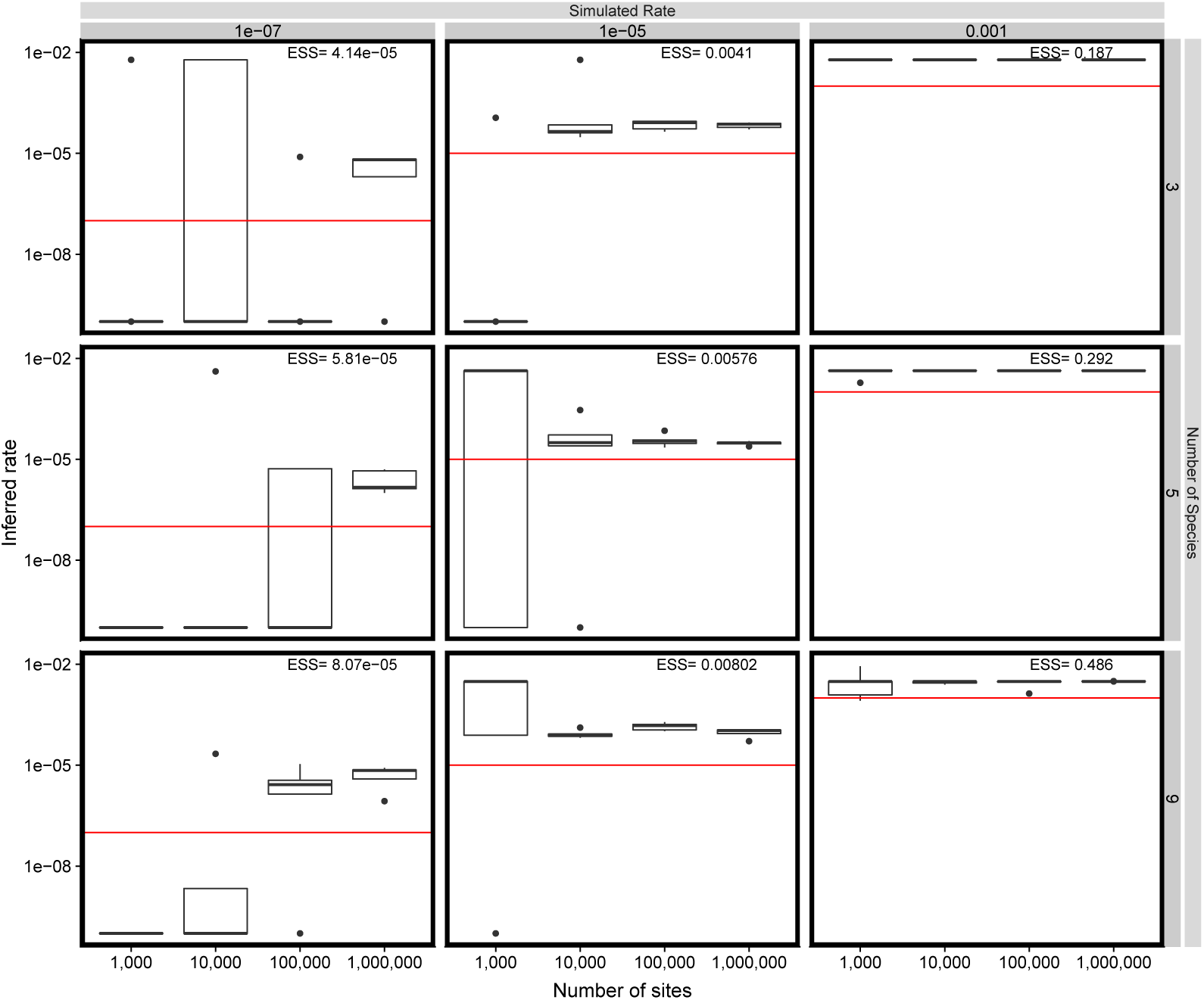
Estimates of γ depend on the number of expected substitutions per site. Raw count data was simulated under the epiPhyloHMM model, with all possible states being enumerated, for genomic regions of varying size (1,000 sites = 250KB) at varying rates for fixed ρ_1_ and π*_A_*. epiPhyloHMM converges to inflated estimates of the correct values given increasing amounts of data, with convergence occurring more rapidly for scenarios with a greater number of expected substitutions per site.

**Figure S3:**
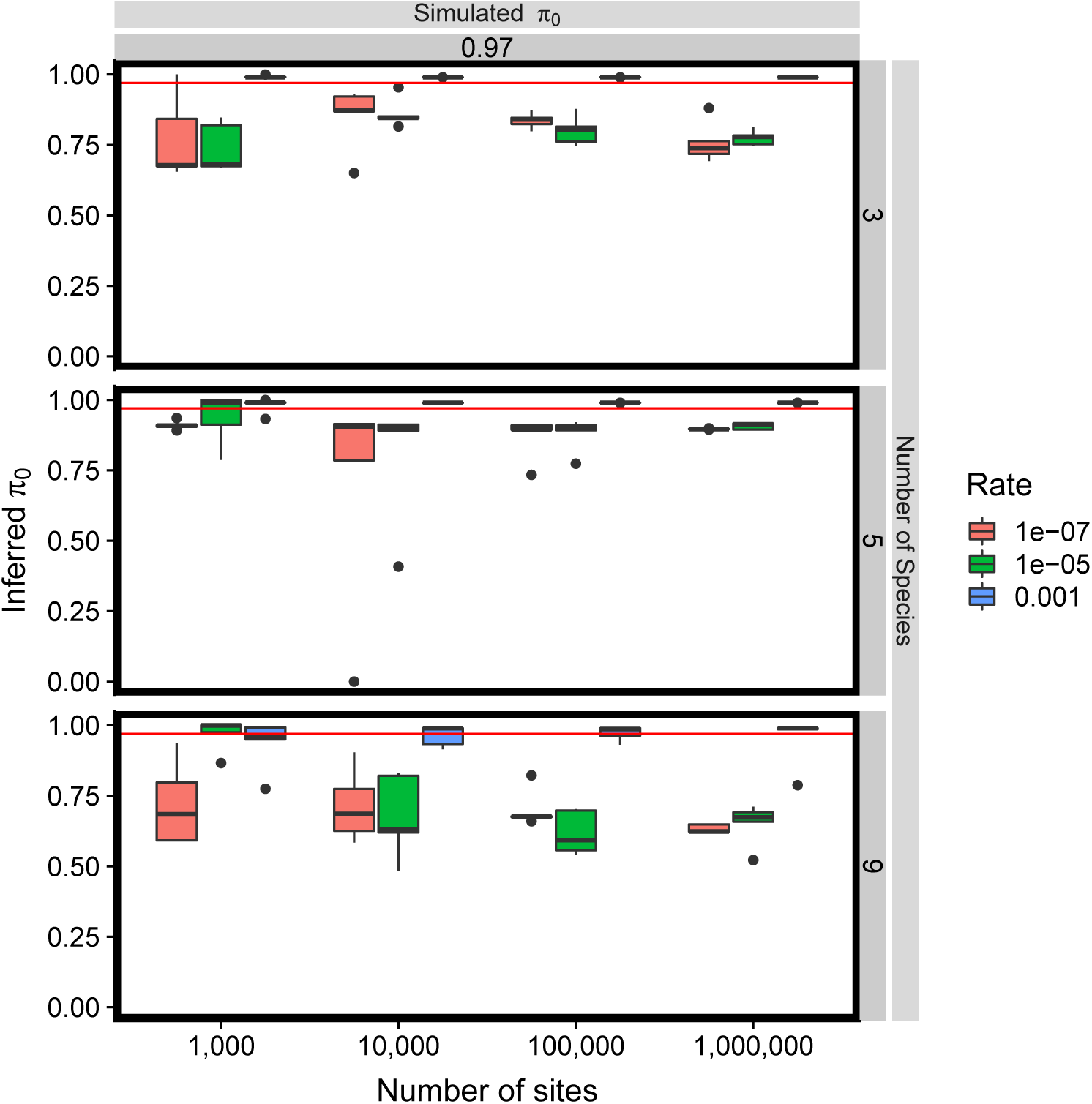
Estimates of π are biased by mis-estimates of rate. Raw count data was simulated under the epiPhyloHMM model, with all possible states being enumerated, for genomic regions of varying size (1,000 sites = 250KB) at varying rates for fixed ρ_1_ and π*_A_*. epiPhyloHMM converges to biased estimates of the correct values depending on the rate.

**Figure S4:**
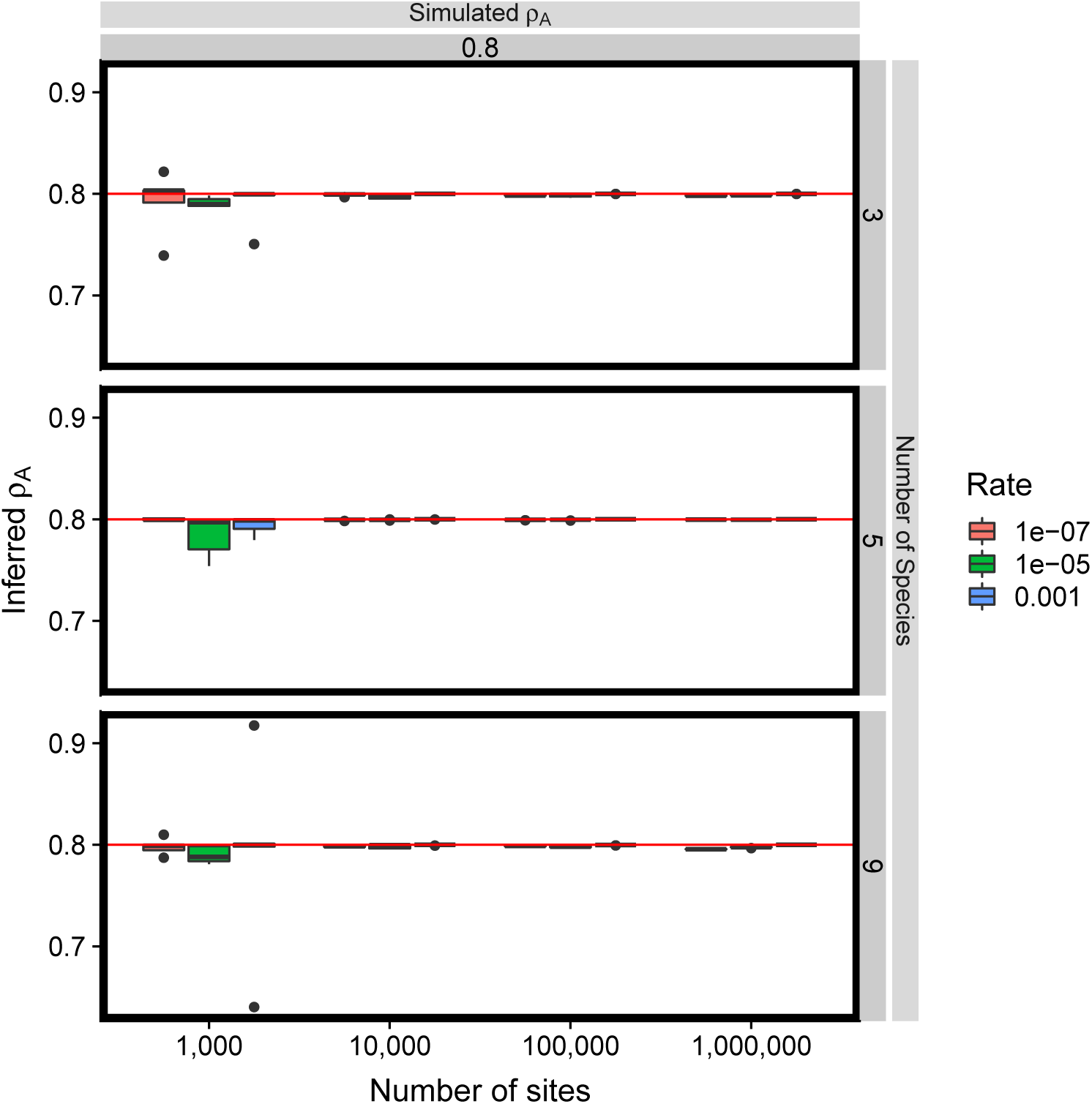
**Estimates of ρ_1_ converge rapidly to the true value R**aw count data was simulated under the epiPhyloHMM model, with all possible states being enumerated, for genomic regions of varying size (1,000 sites = 250KB) at varying rates for fixed ρ_1_ and π*_A_*. epiPhyloHMM converges to correct estimates of the true value of ρ_1_.

**Figure S5:**
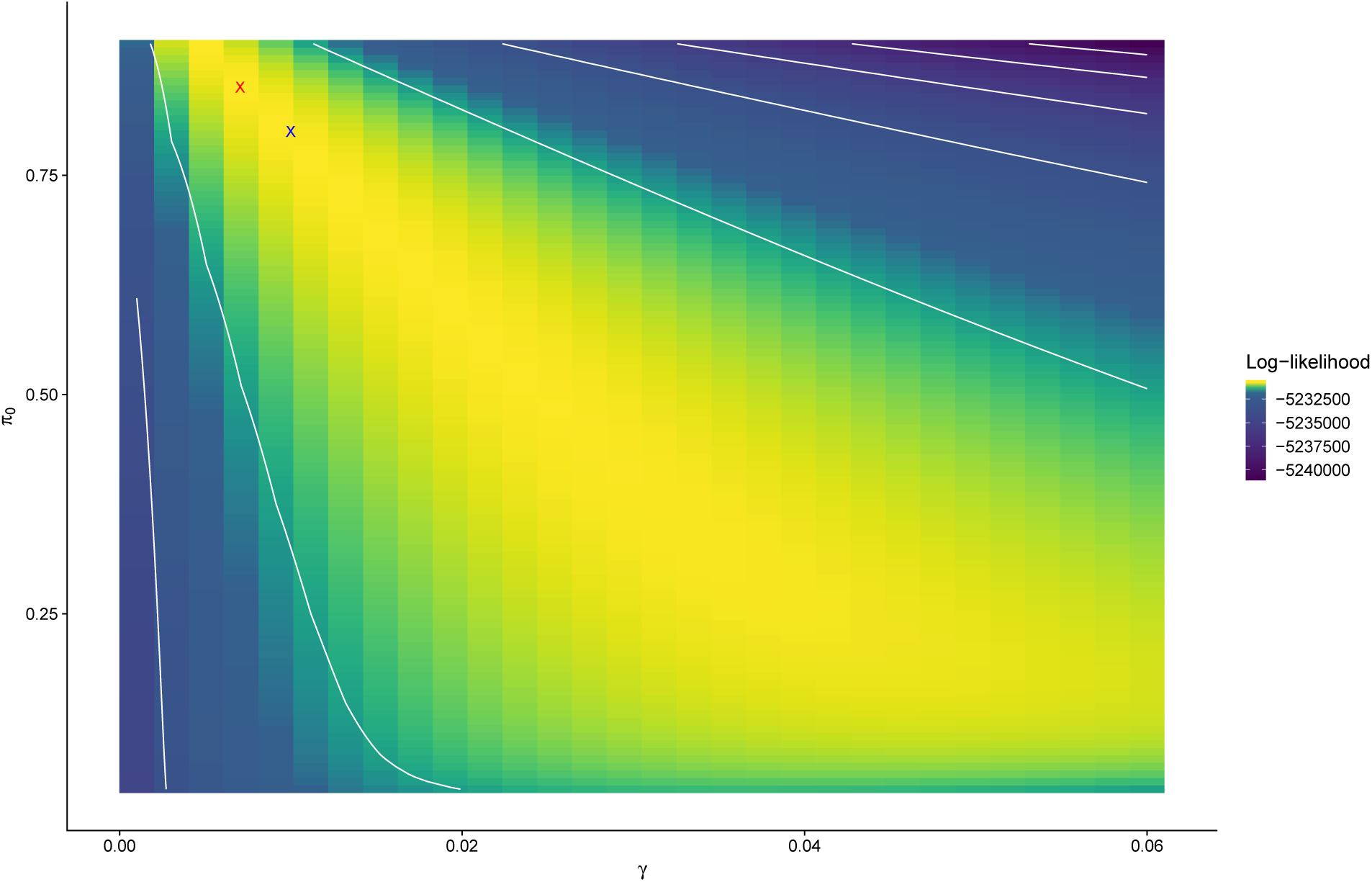
Example of ridge in log-likelihood surface for fitting epiPhylo model. Loglikelihood is computed on finite grid of γ and π values with the auto-correlation parameters (ρ_0_, ρ_1_) fixed to the true values of the simulated data. The blue “X” indicates the true value while the red “X” indicates the MLE parameter values on the computed landscape.

**Figure S6:**
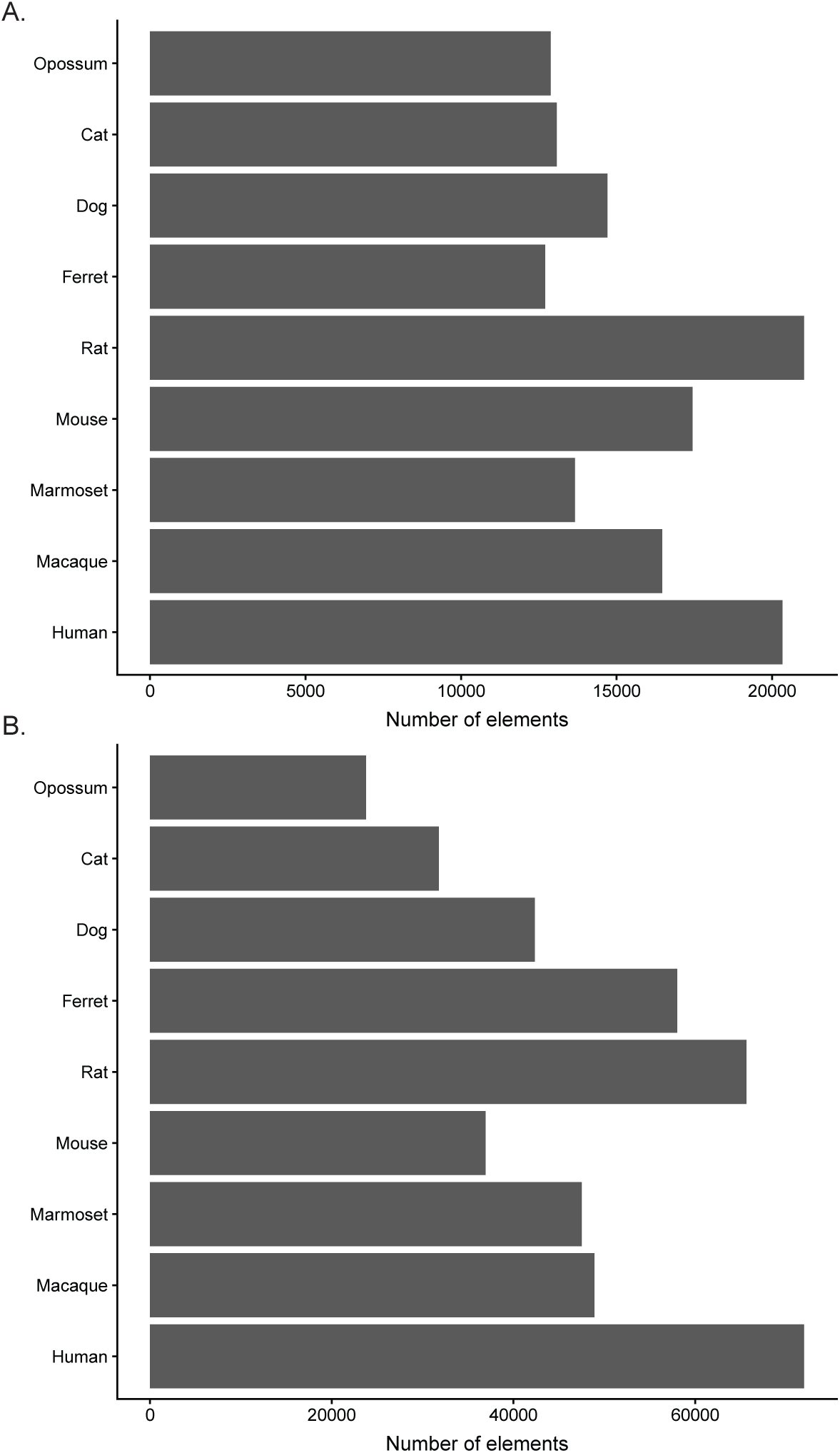
**Number of multi-species epigenetic state calls from epiPhylo based on the Viterbi algorithm** (A) H3K4me3 (B) H3K27ac

**Figure S7:**
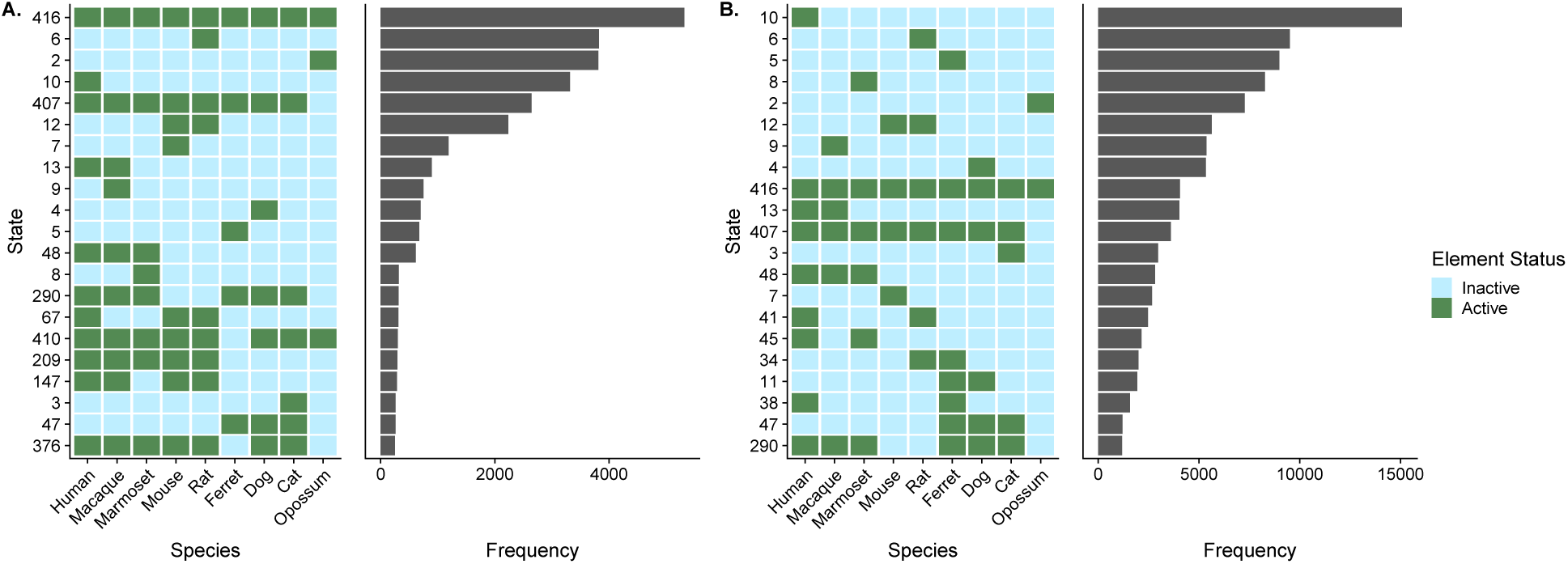
Distribution of common state calls by epiPhyloHMM. (A) The distribution of state calls from epiPhyloHMM for the (A) H3K4me3 mark and (B) the H3K27ac mark. The heatmap describes the configuration of active and inactive elements for each state. All un-visualized states are at lower frequency.

**Figure S8:**
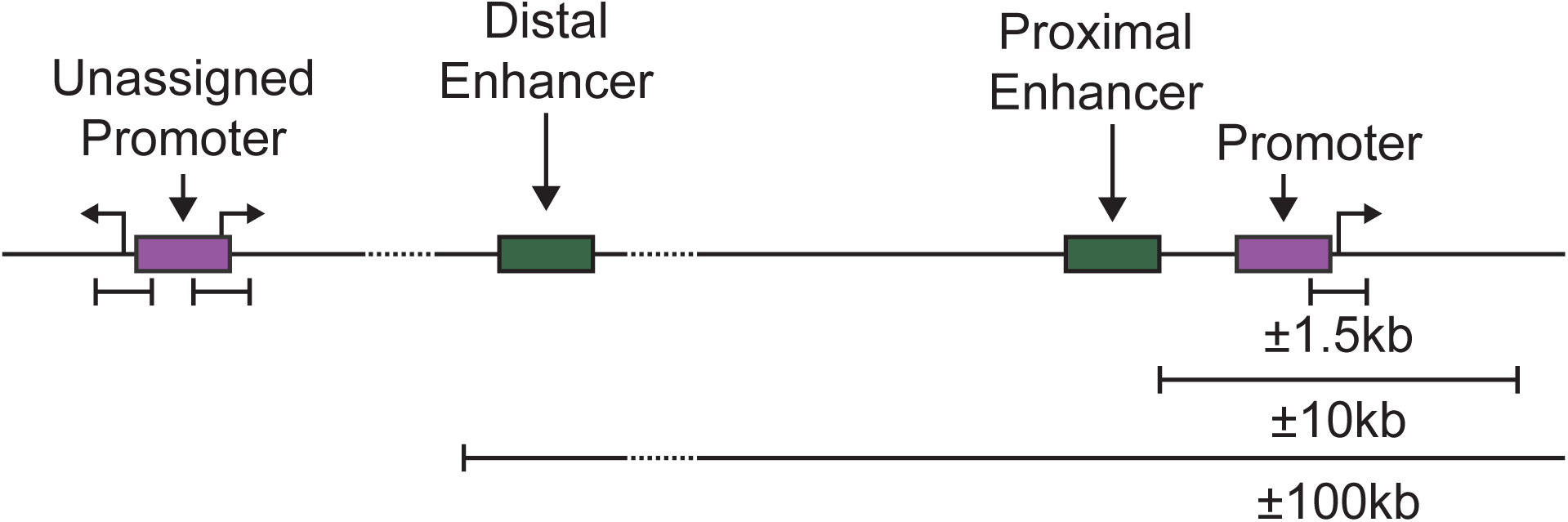
Annotation scheme for epigenetic elements. All genic distances and annotations are based on the human genome build GrCH38^52^. H3K4me3 elements were annotated as promoters if they were within +/ − 1.5kbp of a TSS. If they overlapped with TSS(s) for only one gene, they were associated with that gene. If they overlapped with TSSs from more than one gene, they were annotated as unassigned promoters. The same rules apply for annotating H3K27ac marks as promoters, however there are additional rules for annotating them as enhancers. If a H3K27ac element was within +/−10kbp but not within +/−1.5kbp of a TSS they were annotated as proximal enhancers and assigned analogously to promoters. If an H3K27ac mark was between 10kbp and 100kbp away from the nearest TSS, they were annotated as distal enhancer and assigned to the gene of the nearest TSS. H3K27ac elements further away than 100kbp were annotated as unknown.

**Figure S9:**
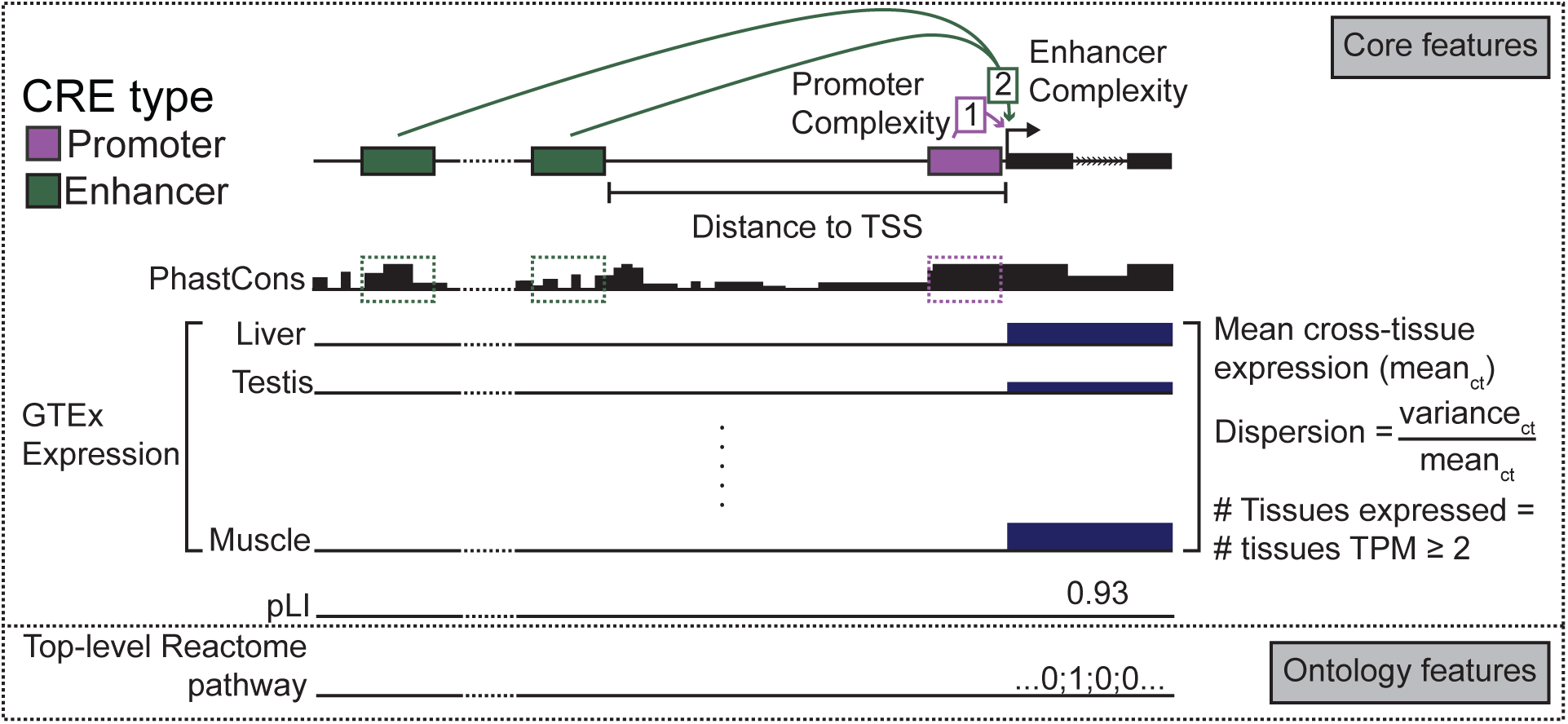
Genomic covariates run with phyloGLM model. Schematic of genomic covariates collected to analyze turnover rates of CREs. Promoter and enhancer complexity is respectively, the number of promoter and enhancers associated with a given gene. Mean phastCons scores are the mean phastCons-100way score in the region covered by a CRE on the human genome (GrCh38). The expression features are computed from GTEx (V6) data assuming independence of samples. Functional catagories are limited to top level Reactome pathways (functional annotations which have no “parent” annotations). It is possible, but very rare for a CRE to have more than one such label.

**Figure S10:**
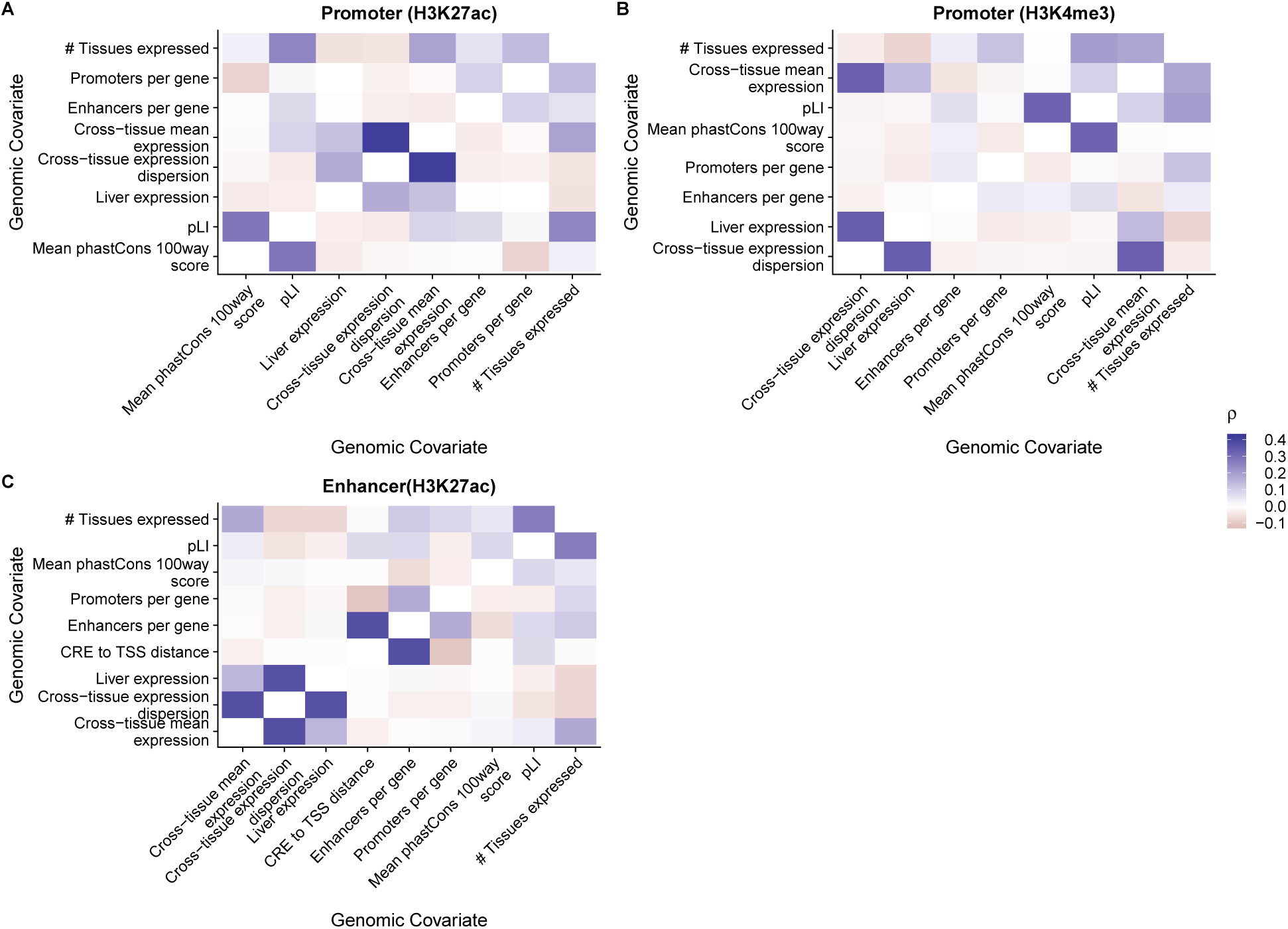
Pearson correlation of genomic covariates for the H3K4me3 and H3K27Ac elements. Pearson correlation of genomic covariates for all elements used to fit a phyloGLM model. Correlations were calculated seperately for (A) H3K27ac promoter, (B) H3K4me3 promoter, and (C) H3K27ac enhancer elements.

**Figure S11:**
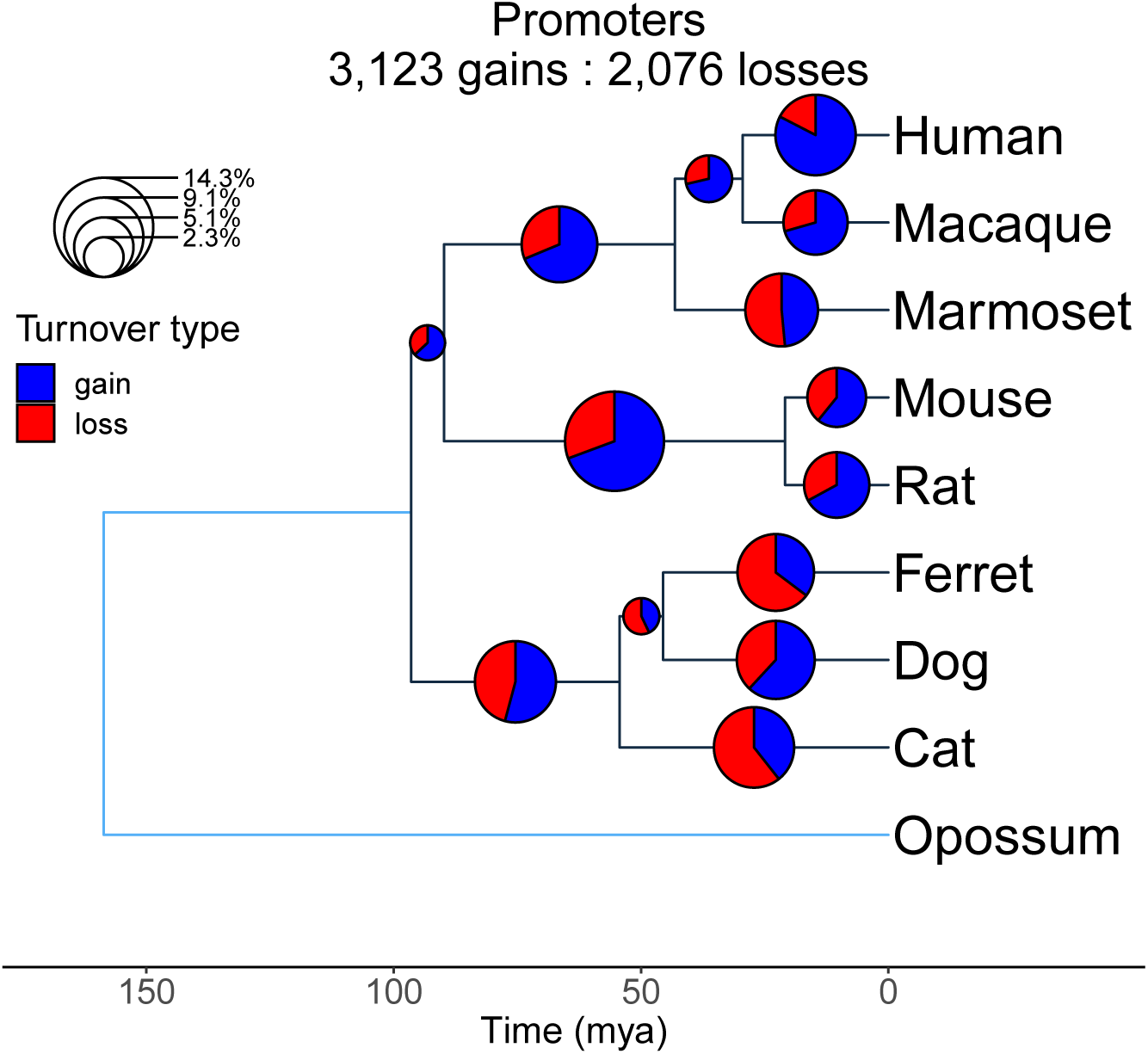
Ancestral reconstruction of gain/loss events on a rooted tree for H3K4me3 elements. Pie chart area per branch is proportional to the fraction of total enhancer/promoter state calls undergoing gain or loss. Numbers of gain/loss events were computed from pairwise marginals of the transition matrices on each branch. Estimates on branches leading to outgroup were removed as there was insufficient information to polarize gain/loss calls.

**Figure S12:**
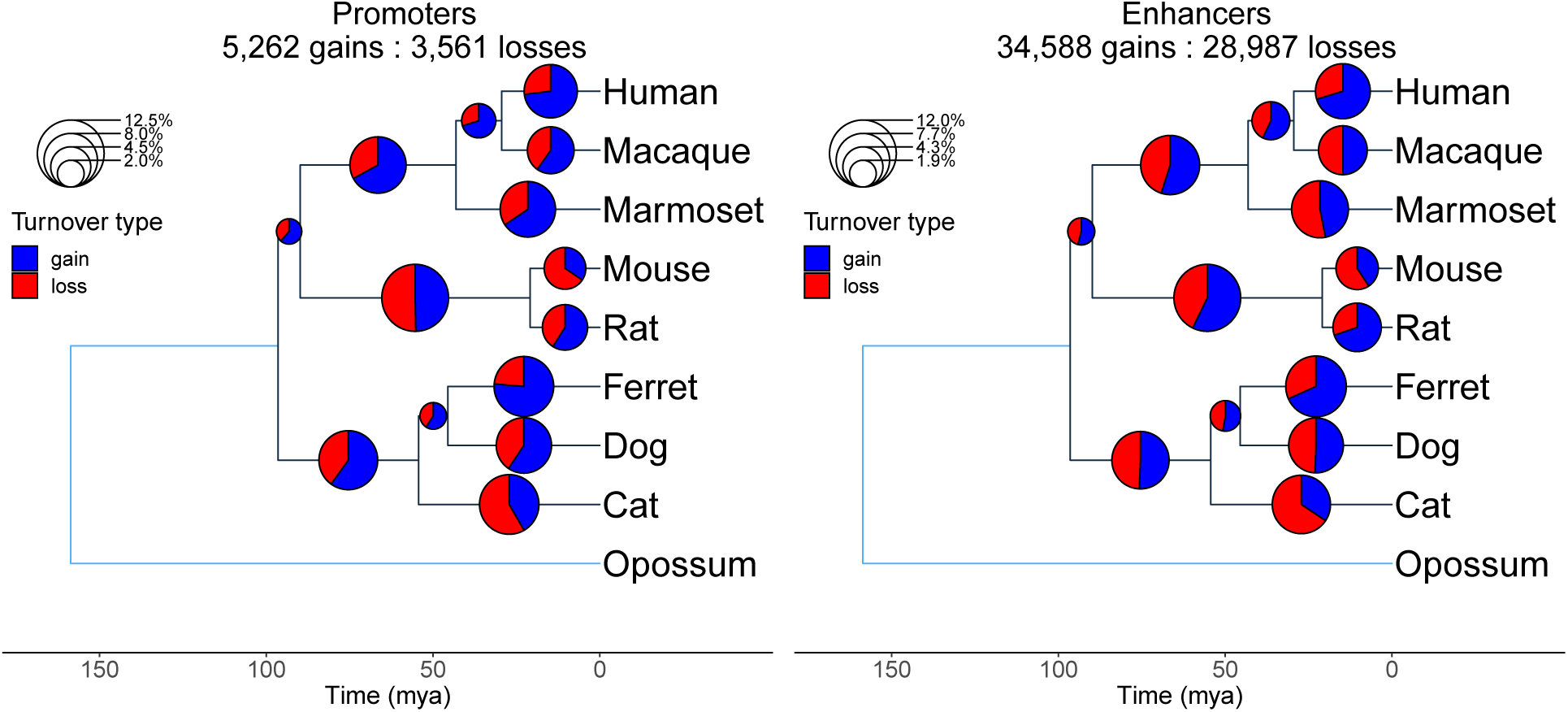
Ancestral reconstruction of gain/loss events on a rooted tree for H3K27ac elements. Pie chart area per branch is proportional to the fraction of total enhancer/promoter state calls undergoing gain or loss. Numbers of gain/loss events were computed from pairwise marginals of the transition matrices on each branch. Estimates on branches leading to outgroup were removed as there was insufficient information to polarize gain/loss calls. Sites with an H3K27ac mark in one or more species were partitioned into enhancers and promoters based on proximity to transcriptional start sites (TSS) in humans (Fig. S8, see **Methods**).

**Figure S13:**
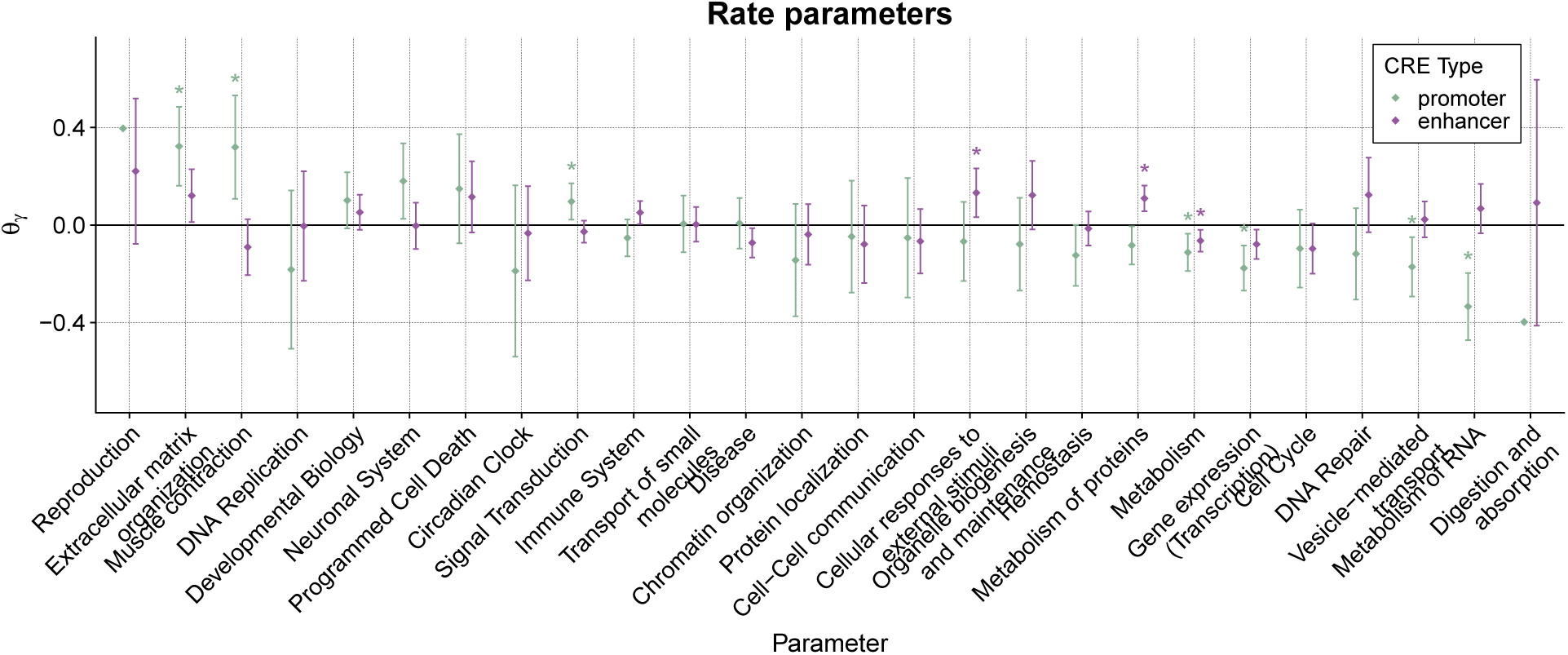
phyloGLM associates some functional characteristics with differential turnover rate. Rate parameter estimates for reactome annotations of genes fitted to H3K27ac data. If parameter is estimated to have a value > 0 is increases the turnover rate; if it is < 0, a decreased turnover rate. Error bars represent approximate 95% confidence intervals derived from the hessian. Parameters with a “*” above them have *FDR*≤ 0.05 calculated via a likelihood ratio test.

